# A Spectral Theory for Wright’s Inbreeding Coefficients and Related Quantities

**DOI:** 10.1101/2020.10.07.329755

**Authors:** Olivier François, Clément Gain

## Abstract

Wright’s inbreeding coefficient, *F*_ST_, is a fundamental measure in population genetics. Assuming a predefined population subdivision, this statistic is classically used to evaluate population structure at a given genomic locus. With large numbers of loci, unsupervised approaches such as principal component analysis (PCA) have, however, become prominent in recent analyses of population structure. In this study, we describe the relationships between Wright’s inbreeding coefficients and PCA for a model of *K* discrete populations. Our theory provides an equivalent definition of *F*_ST_ based on the decomposition of the genotype matrix into between and within-population matrices. The average value of Wright’s *F*_ST_ over all loci included in the genotype matrix can be obtained from the PCA of the between-population matrix. Assuming that a separation condition is fulfilled and for reasonably large data sets, this value of *F*_ST_ approximates the proportion of genetic variation explained by the first (*K* – 1) principal components accurately. The new definition of *F*_ST_ is useful for computing inbreeding coefficients from surrogate genotypes, for example, obtained after correction of experimental artifacts or after removing adaptive genetic variation associated with environmental variables. The relationships between inbreeding coefficients and the spectrum of the genotype matrix not only allow interpretations of PCA results in terms of population genetic concepts but extend those concepts to population genetic analyses accounting for temporal, geographical and environmental contexts.

**Author’s summary:** Principal component analysis (PCA) is the most-frequently used approach to describe population genetic structure from large population genomic data sets. In this study, we show that PCA not only estimates ancestries of sampled individuals, but also computes the average value of Wright’s inbreeding coefficient over the loci included in the genotype matrix. Our result shows that inbreeding coefficients and PCA eigenvalues provide equivalent descriptions of population structure. As a consequence, PCA extends the definition of this coefficient beyond the framework of allelic frequencies. We give examples on how *F*_ST_ can be computed from ancient DNA samples for which genotypes are corrected for coverage, and in an ecological genomic example where a proportion of genetic variation is explained by environmental variables.

## Introduction

Defined by Sewall Wright and Gustave Malécot, the fixation index or inbreeding coefficient, *F*_ST_, measures the amount of genetic diversity found between populations relative to the amount within populations [1, 2]. Used as a measure of population differentiation, *F*_ST_ is among the most widely used descriptive statistics in population and evolutionary genetics [3, 4, 5, 6, 7]. Inbreeding coefficients were originally defined for the analysis of allelic frequencies at a single genetic locus. With the amount of data available to present-day or ancient population genomic studies, principal component analysis (PCA) and model-based estimation algorithms, such as STRUCTURE, have emerged as alternative ways to describe population structure from multilocus genotype matrices [8, 9, 10, 11, 12].

Assuming that the columns of the genotype matrix are either centered or scaled, PCA computes the eigenvalues and eigenvectors of the sample covariance matrix. The first eigenvectors – or axes – summarize the directions which account for most of the genetic variation, and the eigenvalues represent the variances of projected samples along the axes. Eigenvalues and eigenvectors can be computed efficiently by using the singular value decomposition (SVD) of the column-centered data matrix [13]. PCA has been considered very early in human biology, and has become a popular method to study the genetic structure of populations [14, 15]. Inference from PCA is justified from the fact that, similar to STRUCTURE, the projections of individuals on principal axes reveal their degree of admixture with source populations when these sources are represented in the sample [10, 16, 17, 18].

Although the relationships between PCA projections and admixture estimates are well-understood, difficulties of interpreting PCA eigenvalues still remain. The main contributions in that direction were restricted to models of divergence between two populations. The arguments were based on random matrix theory (RMT)[10, 19] and coalescent theory [16]. We note that connections between *F*_ST_ and PCA are not only important for description of population structure, but also in genome scans for selection where the distribution of PCA loadings can be used to detect regions with signature of divergent selection [20, 21, 22, 23]. Based on RMT, Ref. [10] proposed a threshold value of *F*_ST_ for two populations with equal sample sizes [10]. Below the threshold, there should be essentially no evidence of population structure. The coalescent approach relied on a relationship between *F*_ST_ and coalescent time for a pair of genes from a single subpopulation and that of a pair of genes from the collection of subpopulations [6]. For a model of divergence between two populations, theoretical results for coalescent times were used to demonstrate a link between the leading eigenvalue of PCA and *F*_ST_ [16]. Results in Ref.[16] might be extended to simple models of population structure with explicit formulas for coalescent times [24], but the results are not straightforward. While coalescent theory and RMT have provided relationships between *F*_ST_ and PCA in simple cases, the general conditions under which they are valid and their extensions to more than two populations are unknown.

In this study, we develop a spectral theory of genotype matrices to investigate the relationships between PCA and Wright’s coefficients in discrete population models. Our theoretical framework assumes that the observed genotypes correspond to the sampling of *K* discrete populations. Decomposing the genotype matrix as a sum of between and within-population matrices, we extend the results obtained in [10, 16, 19, 25]. Our main result states that the mean value of *F*_ST_ over loci is equal to the squared Hilbert-Schmidt norm of the between-population matrix, which can be computed by a spectral analysis. Under a separation condition bearing on the between and within-population matrices, the sum of the first (*K* – 1) eigenvalues of scaled PCA approximates the mean value of *F*_ST_ over loci. To describe residual variation not explained by the discrete population model, we rely on approximations of the eigenvalues of the within-population matrix from RMT [10, 26].

A corollary of the theory is an alternative definition of inbreeding coefficients that allows us to extend *F*_ST_ to adjusted or *surrogate* genotypes, such as genotype likelihoods and other modifications of allele counts [27]. To illustrate the new definition, we compute *F*_ST_ for ancient human DNA samples after performing correction for genomic coverage and for distortions due to difference in sample ages [28]. In a second application, we compute *F*_ST_ for Scandinavian samples of *Arabidopsis thaliana* after removing genetic variation associated with environmental variables taken from a climate database [29, 30].

## Results and Discussion

### Partitioning of genetic variation

Consider a sample of *n* unrelated individuals for which a large number of loci are genotyped, resulting in a matrix, **X** = (*x_iℓ_*), with *n* rows and *L* columns. For haploids, we set *x_iℓ_* = 0,1, and for diploids *x_iℓ_* = 0,1,2 to count the number of derived alleles at locus *ℓ* for individual *i*. Dealing with autosomes, we simplify our presentation by considering a sample of diploids as being represented by a sample of haploids having twice the original sample size. For unphased data, we take a random phase. Although not a necessary condition, the loci are assumed to be unlinked, or obtained after a linkage disequilibrium (LD)-pruning algorithm applied to the genotype matrix [20, 31]. We use the term locus as a shorthand for single-nucleotide polymorphism (SNP), although most of our analyses could include non-polymorphic sites. Following Wright’s approach to the description of population structure, our main assumption is that individuals are sampled from *K* predefined discrete populations. Examples of discrete population models underlying our assumptions include Wright’s island models, coalescent models of divergence and *F*-models [6, 32, 33]. Application to *F*-models will be described afterwards.

To analyze population structure, PCA can be performed after centering and some-times after scaling the genotype matrix. The scaled matrix is denoted by **Z**^sc^ and the unscaled centered matrix is denoted by **Z**. Scaled PCA computes the eigenvalues, 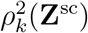, of the empirical correlation matrix. Unscaled PCA computes the eigenvalues, 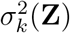, of the empirical covariance matrix [9, 26]. The eigenvalues are ranked in decreasing order, and 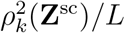 is usually interpreted as the proportion of variance explained by the *k*th axis of the PCA. PCA can be performed via the SVD algorithm. In this case, the eigenvalues of scaled (or unscaled) PCA correspond to the squared singular values of the scaled (or centered) matrix divided by 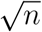 [9, 26].

To establish relationships between PCA and inbreeding coefficients, we decompose the centered matrix into a sum of two matrices, **Z** = **Z**_ST_ + **Z**_S_, corresponding to between and within-population components. The decomposition is performed as follows. Let *i* be an individual sampled from population *k*. At a particular locus, *ℓ*, the genotype, *x_iℓ_*, is equal 0 or 1 (derived allele), and *p_kℓ_* denotes the derived allele frequency in population *k* at this locus. The coefficient of the centered matrix, *z_iℓ_*,s is equal to 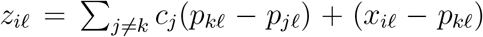, where *c_k_* = *n_k_/n* represents the proportion of individuals sampled from population *k*. In this formulation, the between-population matrix, **Z**_ST_, has general term 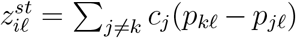, repeated for all individuals in population *k*. By construction, the rank of **Z**_ST_ is equal to (*K* – 1). The within-population matrix, **Z**_S_, has general term 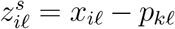. A very similar decomposition holds for the scaled matrix, defined as 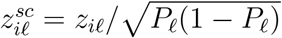, where 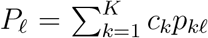 is the derived allele frequency in the total sample at locus *ℓ* (see Box 1 for the notations).

### Spectral analysis of inbreeding coefficients

Consider *n* samples from *K* discrete populations and define *D*_ST_ and *F*_ST_ according to Wright’s [1] and Nei’s [4, 34] equations, allowing for unequal population sample sizes. At a particular locus, set 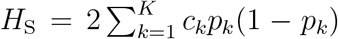 and *H*_T_ = 2*P*(1 – *P*), then we have *D*_ST_ = *H*_T_ – *H*_S_. Wright’s inbreeding coefficient is defined as

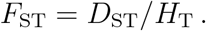

Our main result states that the mean value of *F*_ST_ across loci can be computed from the singular values of the between-population matrix, 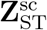. A similar relationship is also established for *D*_ST_ and for the unscaled matrix, **Z**_ST_. The singular values of the between and within-population matrices can be evaluated from the SVD algorithm. The computational cost of those operations is equal to the computational cost of the PCA of the genotype matrix, of order *O*(*n*^2^ *L*). This cost could be reduced by using various methods, for example by computing the first *K* – 1 singular values only. All conclusions below are valid regardless of any population genetic model. The results also remain valid when genotypes are conditioned on having minor allele frequency greater than a given threshold, and when the loci are physically linked or when they exhibit LD.

#### Theorem 1.

*Let K* ≥ 2. *Let* **Z** *and* **Z**^sc^ *be the unscaled and scaled genotype matrix respectively. Define* **Z**_ST_ *and* 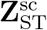 *as in the previous section. We have*

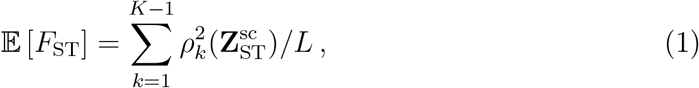

*and*

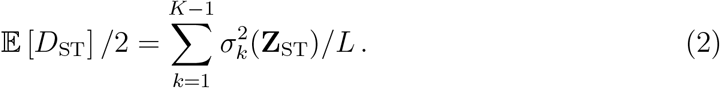

The key arguments for those results involve matrix norms, and they can be found in *Supplementary Information*. By the Pythagorean theorem, we have ∥*Z*∥^2^ = ∥**Z**_ST_∥^2^ + ∥**Z**_S_^2^∥ (*Supplementary Information*). According to this result, Theorem 1 can be reformulated using within-population matrices as follows.

#### Corollary.

*Let K* ≥ 2. *Let* **Z** *and* **Z**^sc^ *be the unscaled and scaled genotype matrix respectively. Define* **Z**_S_ *and* 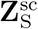 *as in the previous section. We have*

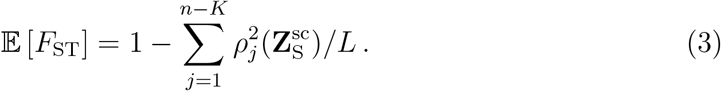

*and*

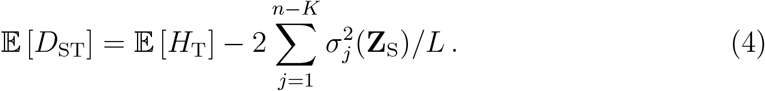

### Inbreeding coefficients for surrogate genotypes

Besides an interest in connecting population genetic theory to the spectrum of the genotype matrix, the results in equations (1) and (2) have important consequences for data analysis. First, the ? results support the definition of *F*_ST_ for multilocus genotypes as an average of ratios ? rather than a ratio of averages [35, 36]. More importantly, Theorem 1 leads to alter; native definitions of Nei’s *D*_ST_ and Wright’s *F*_ST_ from population genetic data. As will be demonstrated by applications to ancient DNA and to ecological genomics, the main interest in those new definitions is their straightforward extension to adjusted genotypic matrices, providing statistics analogous to *F*_ST_ for modified genotypic values. For example, adjusted genotypic matrices arise when correcting for biases due to technical artifacts, including batch effects and genomic coverage in population genomic data [37, 38]. In general, *F*_ST_ could be adjusted for any specific effect by; considering the residuals of latent factor regression models [39, 40, 41]. More specifically, for **Z** (or **Z**^sc^) and for a set of measured covariates, **Y**, latent factor regression models estimate a matrix of surrogate genotypes, **W**, by adjusting a regression model of the form

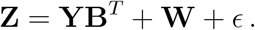

In this model, the **B** matrix contains effect sizes for each variable in **Y**, and *ϵ* is a matrix that represents centered errors. The latent matrix **W** has a specified rank, *k*, lower than *n* minus the number of covariates. The rank *k* corresponds to the number of latent factors incorporated in the model. The matrix **Z**^adj^ = **W** + *ϵ* leads to a definition of an adjusted inbreeding coefficient, 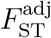. The adjusted inbreeding coefficient can be computed as the squared norm of the between-population matrix, 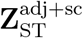, after scaling. Note that this definition considers quantitative values observed at each locus, and is similar to a population genetic quantity called Q_ST_ [42, 43]. Alternatively, 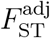 can be computed from the average coefficient of determination, *R*^2^, obtained from the regression of each scaled surrogate genotype on the population labels. The definitions are equivalent, and we have

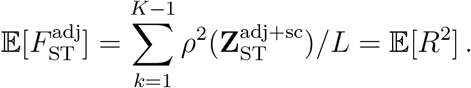

### Inbreeding coefficients and PCA eigenvalues

Having established that the mean value of *F*_ST_ across loci can be computed from the leading eigenvalues of the between-population matrix, 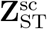, the next question is to ask whether similar results hold for the leading eigenvalues of the PCA of the scaled matrix,

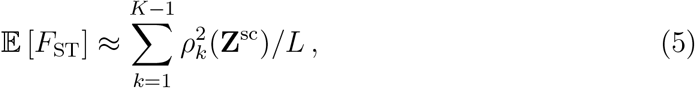

and for the PCA of the centered matrix,

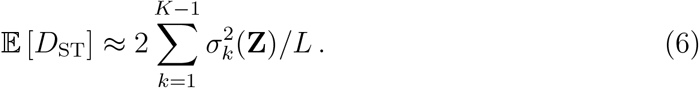

Those results require that the ranked eigenvalues of 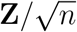 sort into approximate eigenvalues of 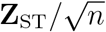 followed by approximate eigenvalues of 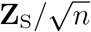. Said differently, the (*K* – 1)th singular value of 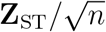 must separate from the leading singular value of 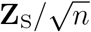

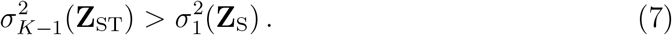

We suppose that the ratio *L*/*n* is constant for large *L* and *n*, and make the following assumptions: 1) The separation condition (6) is verified, 2) The leading eigenvalue of 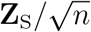 is of order 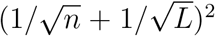 (RMT hypothesis). Then, under those conditions, the accuracy of the approximation in equations (5) and (6) is of order *O*(*K*/*L*). More precisely, for any singular value, *σ_k_*(**Z**_ST_), of 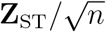, there exists a singular value, *σ_k_*(**Z**), of 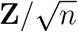 such that we have

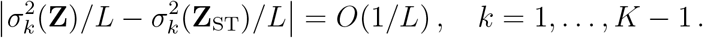

A similar result holds for the first *K* – 1 eigenvalues of scaled matrices, 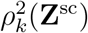 and 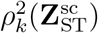. In other words, the mean value of *F*_ST_ across loci can be approximated from the sum of the (*K* – 1) leading eigenvalues of the PCA with an accuracy proportional to the number of populations and to the inverse of the number of unlinked loci in the genotype matrix. Mathematical arguments for those results are detailed in *Supplementary Information*.

Poor approximations may be caused by insufficient sample size, incorrect definition of populations, inclusion of individuals with mixed ancestry, spatial structure, etc. Poor approximations may also be accompanied by failure to verify the separation condition (7), as our simulations will illustrate afterwards. In addition, the results show that *F*_ST_ and PCA exhibit similar biases, for example, when the sampling design is uneven or when loci are filtered out of the genotype matrix [16, 31, 44]. We note that the RMT hypothesis for the residual matrix, **Z**_S_, is difficult to prove for population genetic models. Like [10], we will rely on simulations to show that RMT describes residual variation in population genetic models accurately. In empirical data analyses, checking the residual matrix for agreement with RMT will also provide an informal test for the number of components in PCA similar to Cattel’s elbow rule [45].

### Brief example

To illustrate the approximation of 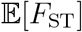 by the leading eigen-values of the PCA, we present a short simulation example, in which a genotype matrix was generated according to a three-population *F*-model. Simulation studies of *F*-models and additional examples based on real data will be developed more extensively later on. In this first simulation example, the average ancestral frequency was equal to 20%, and the drift parameters for the three populations were equal to *F*_1_ = 5%, *F*_2_ = 10% and *F*_3_ = 30%. Populations 1 and 2 were genetically closer to each other than to population 3, which was the most diverged population. Three hundred samples (*n_k_* = 100, for *k* = 1, 2, 3) were genotyped at 10,000 loci. PCA was conducted on *L* = 9740 SNP loci after monomorphic loci were removed. The average value of *F*_ST_ across loci was equal to 9.52%, and approximated the sum of the leading eigenvalues of the PCA (9.54%) accurately. The first axes of the PCA explained 6.78%, 2.76%, and 0.47% of the total variation respectively (Figure 1). The non-null eigenvalues of 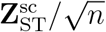, 6.77% and 2.75%, were close to the values obtained for the first two PCs. As stated in Theorem 1, their sum was equal to 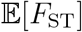. Clearly, the smallest eigenvalue of 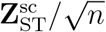 separated from the leading value of 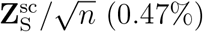, which was close to the eigenvalue for the third PC and to its prediction from RMT (0.31%, Figure 1).

**Figure 1.**
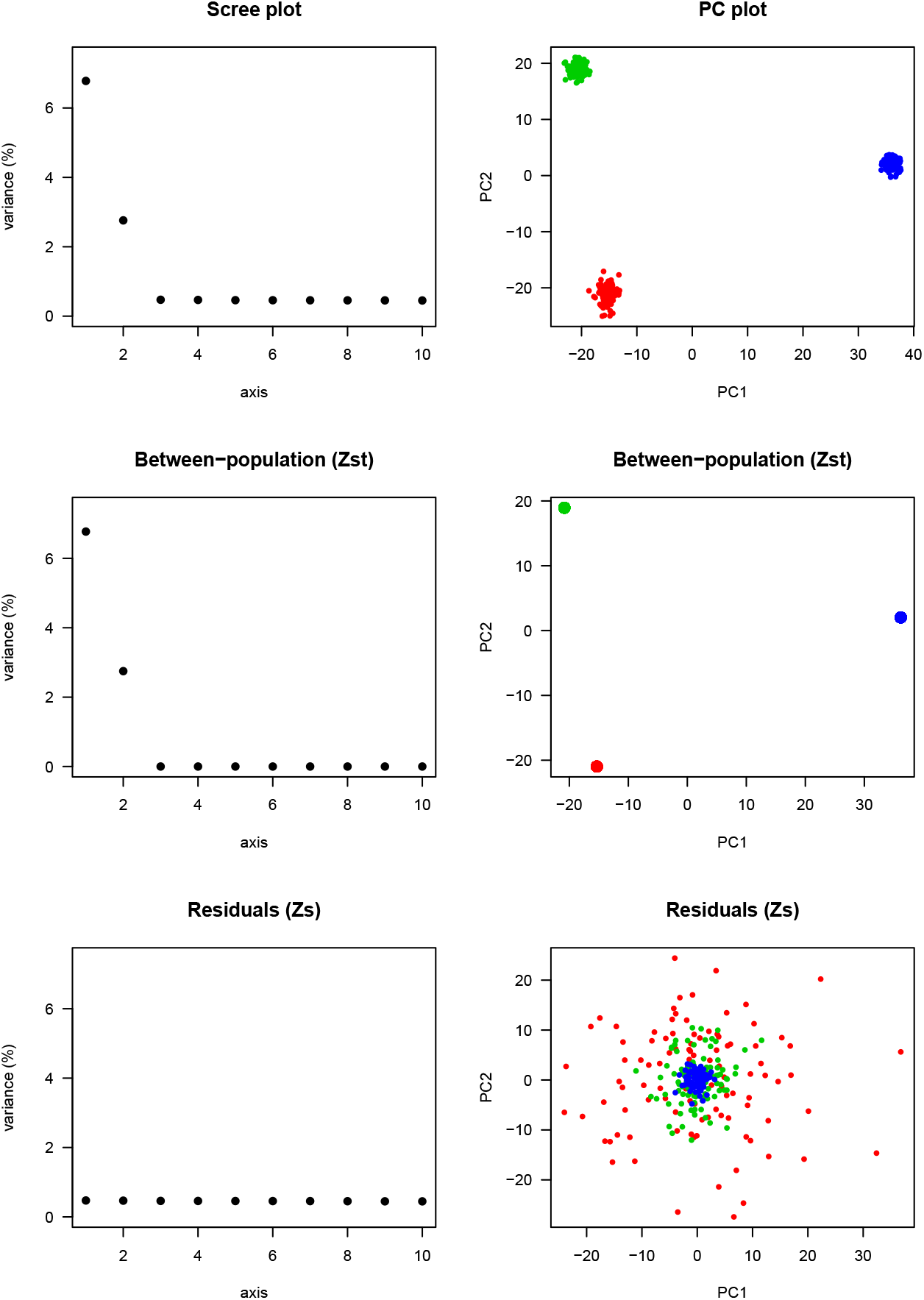
Spectral analysis of a three-population model. PCA scree plots and PC plots for the scaled matrix, **Z**^sc^, for the between-population matrix, 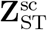, and for the residual matrix, 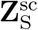 of simulated data. The simulation was performed for *n* = 300 individuals and an *F*-model (*F*_1_ = 5%, *F*_2_ = 10%, *F*_3_ = 30%) with ancestral frequency drawn from a beta(1,4) distribution.

To show the effect of having incorrectly labelled population samples, we considered the same genotype matrix, and replicated the analyses after grouping the “paraphyletic” samples from populations 2 and 3 against the least diverged sample from population 1 (Figure 2, *n*_1_ = 200 and *n*_2_ = 100, *K* = 2). The average *F*_ST_ value was equal to 3.46%, and failed to approximate the first eigenvalue of the PCA (6.78%). The leading value of 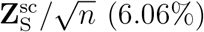 did not verify the separation condition, and it differed from its RMT prediction (0.32%). The PC plot for 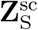 provided evidence of residual population structure within the paraphyletic population sample. Like for a regression analysis, those results outlined the usefulness of visualizing the residual matrix in order to evaluate the number of populations from the genotype matrix (Figure 1 and Figure 2).

**Figure 2.**
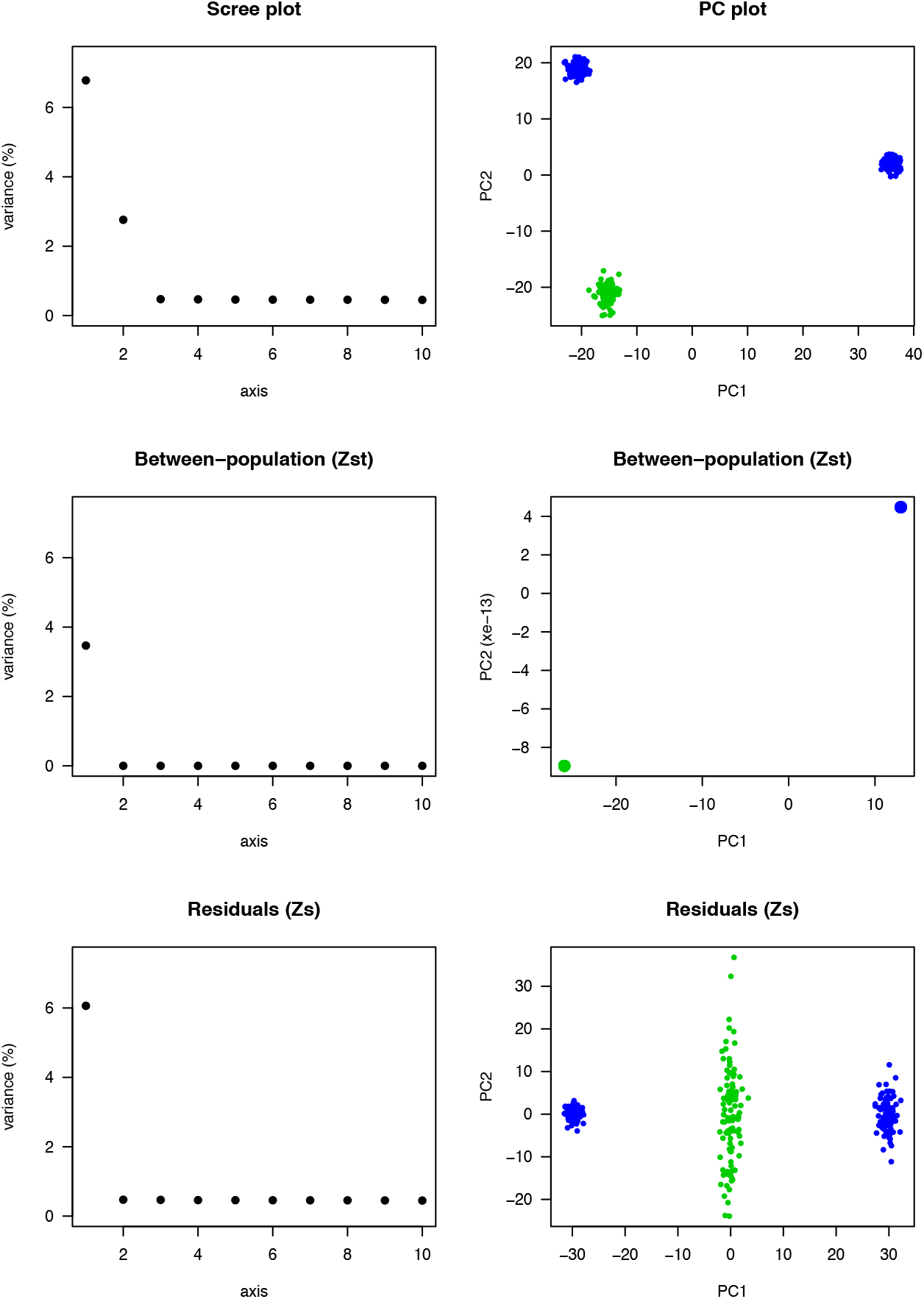
Spectral analysis with incorrectly labelled population samples (*K* = 2). For the same genotype matrix as in Figure 1, samples from populations 2 and 3 (blue) were grouped against population 1 (green). PCA scree plots and PC plots for the scaled matrix, **Z**^sc^, for the between-population matrix, 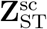, and for the residual matrix, 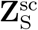 of simulated data. *F*_ST_ was lower than the leading eigenvalue of the residual matrix, and it differed from the first PC eigenvalue.

### Single population models

In a first series of simulations without population structure, we investigated whether RMT predictions were valid for *F*-models. For single population *F*-models, the results supported that the leading eigenvalues of PCA were accurately predicted by the Marchenko-Pastur distribution (Supplementary Figure S1). Then we investigated whether condition (7) could be verified when there was no structure in the data, and two population samples were wrongly defined from a preliminary structure analysis. We ran two-hundred simulations of single population models (*n* = 100 and *L* ≈ 10, 000), and, for each data set, we partitioned the samples in two groups according to the sign of their first principal component. This procedure maximized the likelihood of detecting artificial groups, leading to an average *F*_ST_ ≈ 1.1%. For those artificial groups, we computed the non-null singular value of the between-population matrix, 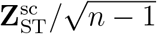, and the leading singular value of the within-population matrix, 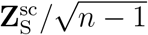. For the simulations, the separation condition was never verified, rejecting population structure in all cases (Supplementary Figure S2A). For smaller sample sizes (*n* = 10 and *L* ≈ 1,000), the separation condition was erroneously checked in 21% simulations, indicating that we had less power to discriminate among artificial groups with small sample sizes (Supplementary Figure S2B). Those results were also consistent with difficulties reported for between-group PCA [46].

### Two-population models

To check whether the expected values of *F*_ST_ and *D*_ST_ were approximated from the first eigenvalues of PCA, we performed simulations of *F*-models with two populations. For these simulations, the separation condition was verified in all data sets. There was an almost perfect fit of the leading eigenvalue for centered PCA, 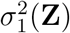, with the average value of *D*_ST_/2 across loci and with its theoretical value in *F*-models (Figure 3A, *Supplementary Information*, Supplementary Figure S3). There was also an almost perfect fit of the leading eigenvalue of scaled PCA, 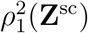, with the average value of *F*_ST_ across loci (Figure 3C). The second largest eigenvalues were accurately predicted by RMT both for unscaled and for scaled PCA (Figure 3 B-D). To detail those results for particular values of drift coefficients, we performed additional simulations for *F*_1_ = *F*_2_ = 7%, also investigating the distribution of eigenvalues of the residual matrix (Figure 4). In a sample of *n* = 200 individuals and *L* ≈ 85, 500 SNPs, the first PC axis explained 3.11% of total genetic variation, corresponding to the average of *F*_ST_ across loci (3.11%, Figure 4A). The separation of between and within-population components was verified, and the second eigenvalue (0.536%) was very close to its prediction from RMT, given by 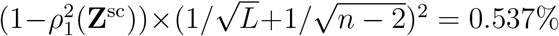 (Figure 4A). The distribution of residual eigenvalues, corresponding to within-population variation, was accurately modelled by the Marchenko-Pastur probability density function (Figure 4B). With a smaller sample of *n* = 20 individuals and *L* ≈ 12,500 SNPs, the leading axis explained 5.24% of the total genetic variation, still matching the value of *F*_ST_ across loci (5.23%, Figure 4C). The Marchenko-Pastur density remained an accurate approximation to the bulk spectrum of residual eigenvalues (Figure 4D).

**Figure 3.**
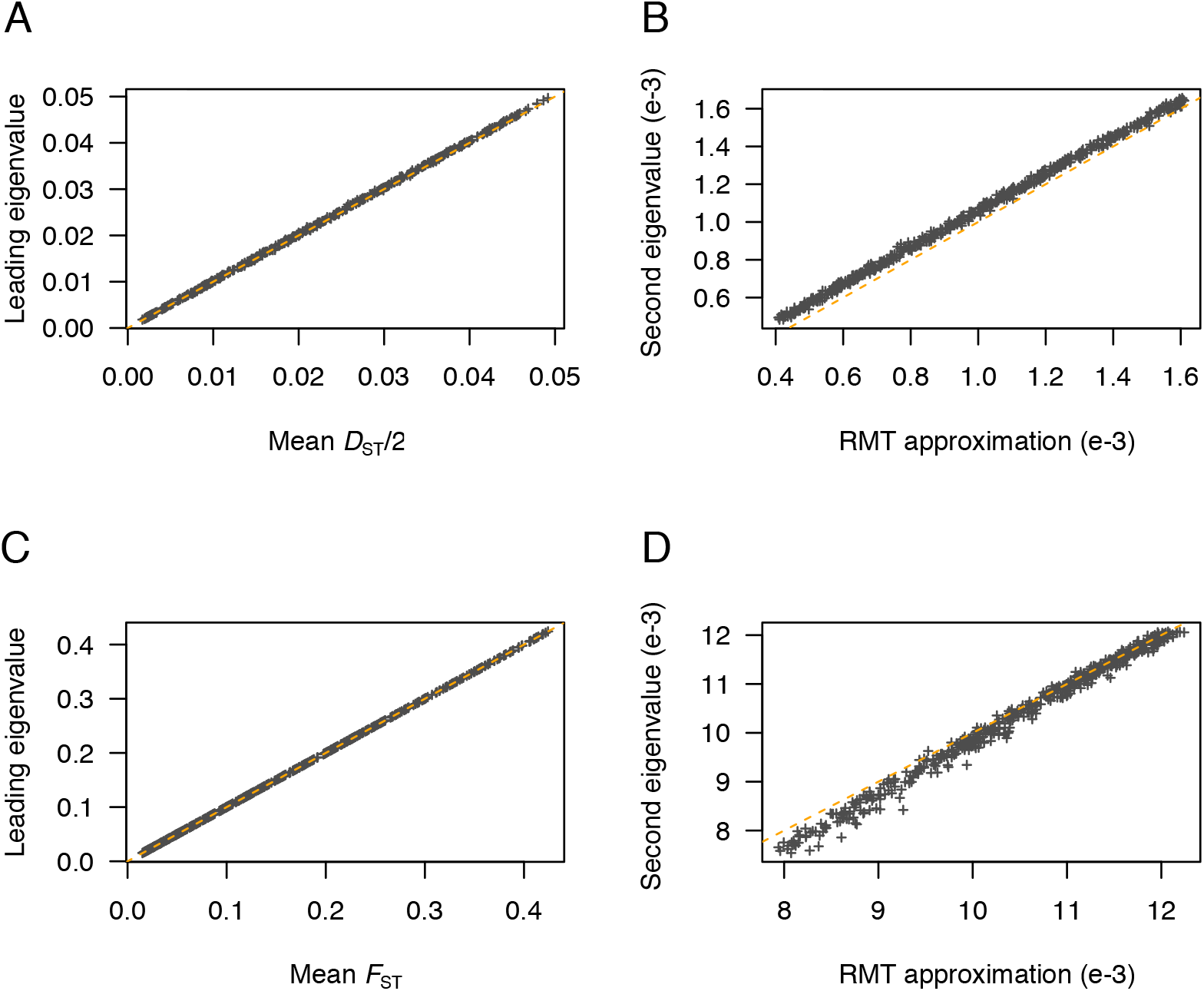
Comparison of *D*_ST_ and *F*_ST_ estimates with the leading PCA eigenvalues in two-population models. **(A)** Leading eigenvalues of centered PCA as a function of the mean of *D*_ST_/2 across loci. **(B)** Second eigenvalue of centered PCA as a function of its approximation from RMT. **(C)** Leading eigenvalues of scaled PCA as a function of the mean of *F*_ST_ across loci. **(D)** Second eigenvalue of scaled PCA as a function of its approximation from RMT, which is given by 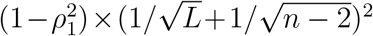 (approximation of the largest eigenvalue of the residual matrix). The dashed lines correspond to the diagonal *y* = *x*. Simulations of *F*-models were performed for *n* = 100 individuals (inbreeding coefficients between 1% and 75%, first population sample proportion between 10% and 50%, ancestral frequency was drawn from a beta(1,4) distribution).

**Figure 4.**
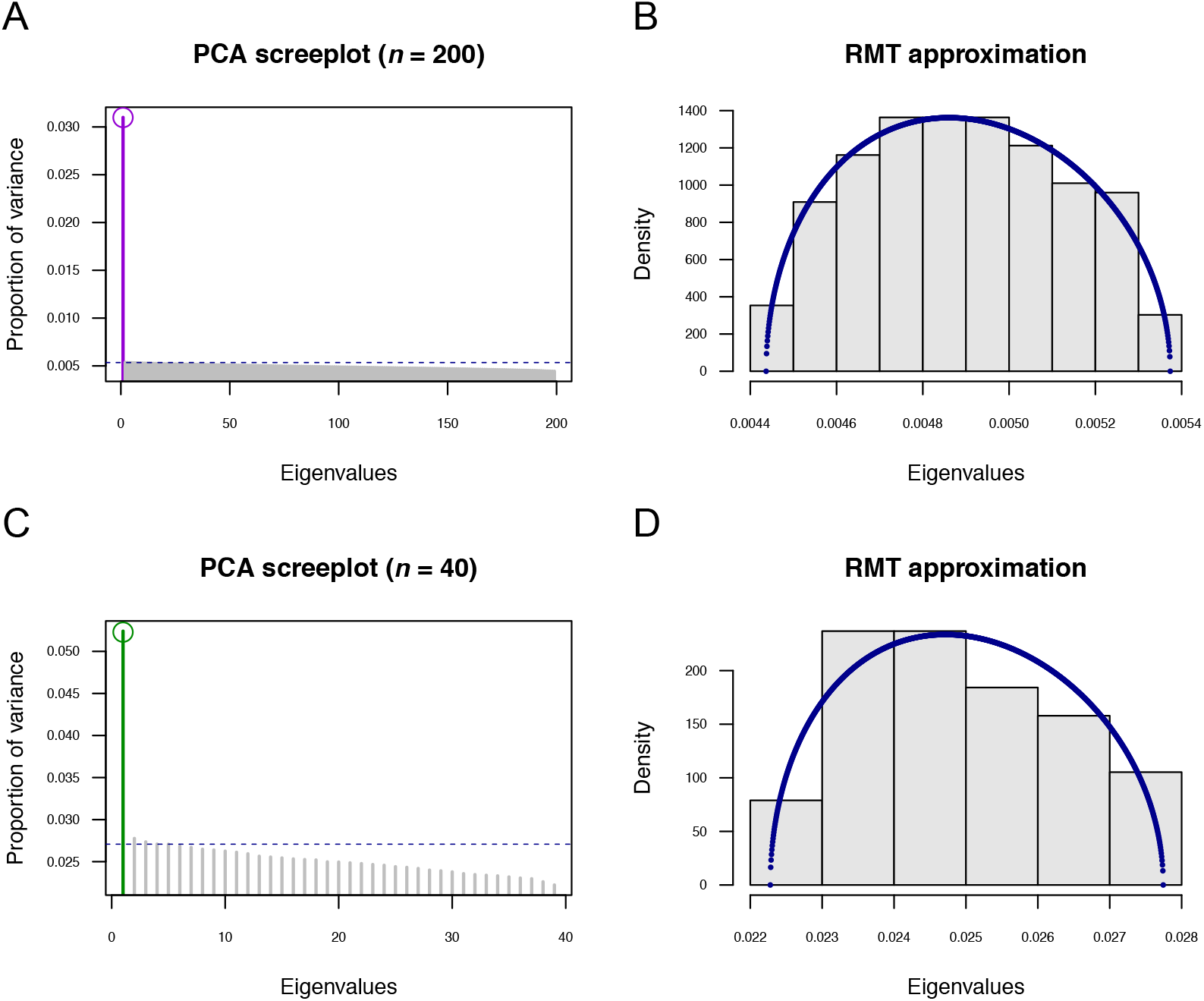
Scree plots and RMT approximations in two-population models. **(A)** Proportion of variance (eigenvalues) explained by PC axes, with a circle symbol representing the mean of *F*_ST_ across loci for *n* = 200 individuals and *L* = 85, 540 SNPs. **(B)** Histogram of eigenvalues of the residual matrix, 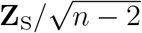, for the data simulated in A. **(C)** Proportion of variance for *n* = 40 individuals and *L* = 12, 650 SNPs. **(D)** Histogram of eigenvalues of the residual matrix for the data simulated in C. The dashed lines in PCA scree-plots represent the RMT approximation of the leading eigenvalue of the residual matrix. The blue curve represents the Marchenko-Pastur probability density. Simulations of *F*-models were performed with *p*_anc_ drawn from a beta(1,9) distribution and *F*_1_ = *F*_2_ = 7%.

In another series of simulations with *F*_1_ = *F*_2_ = 2%, we evaluated whether the sum of the leading eigenvalues for scaled PCA was close to the average of *F*_ST_ across loci when the separation condition was verified (Supplementary Figure S4). For small sample sizes (*n* = 20 and *L* = 100), the separation condition was not verified and the approximation was poor. For intermediate sample sizes (*n* = 60 and *L* = 1000), the approximation was more accurate when the separation condition was verified than when it was not. For large sample sizes (*n* = 100 and *L* = 10000), the separation condition was always verified and the approximation was accurate. Although the *L/n* ratio was not kept to a constant value, the accuracy of the approximation agreed with a predicted order of *O*(1/*L*) (Supplementary Figure S5).

Next, we studied the relationship between leading eigenvalues and sample size, for *L* = 100 loci and *L* = 100, 000 loci (Supplementary Figure S6). For smaller number of loci (*L* = 100) and smaller samples (*n* ≤ 80), the data failed to verify the separation condition in some simulations. In those cases, population structure was not correctly evaluated by 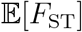. The separation condition was verified in about 35% cases for *n* = 10 and in about 95% cases for *n* ≈ 80. For the larger number of loci (*L* ≥ 100000) or for larger sample sizes (*n* ≥ 100), the separation condition was verified in all cases, and the leading eigenvalue converged to the theoretical value of 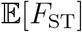 for infinitely large sample sizes. As for between-group PCA, the results suggest exaggerated differences among groups when sample sizes are very small relative to the number of loci [47].

### More than three-population models

For *F*-models, the eigenvalues of the theoretical covariance matrix were analysed formally for small numbers of populations (*Supplementary Information*). To achieve this goal, we considered the co-variance matrix of the random vector **z** defined by 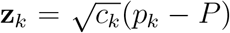, for all *k* = 1 to *K*. The *K* × *K* covariance matrix of the random vector **z** could be obtained from the drift statistics 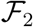 and 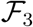 defined in [48, 49]. The coefficients of this matrix are 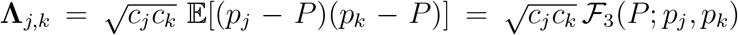, for *j* ≠ *k*, and 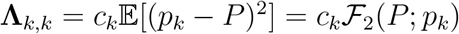 otherwise. For *K* = 3, the eigenvalues of the Λ matrix were computed exactly (*Supplementary Information*).

We performed simulations of three-population *F*-models to check whether the data agreed with theoretical predictions for the leading eigenvalues, λ_1_, and for *D*_ST_ and *F*_ST_. With random drift coefficients (*n* = 100, *L* = 20000), the separation condition was verified in all simulated data sets. An almost perfect agreement between λ_1_ + λ_2_ and the mean value of *D*_ST_/2 (unscaled PCA) or *F*_ST_ (scaled PCA) was observed (Supplementary Figure S7 AC). The leading eigenvalue of unscaled PCA exhibited a small but visible bias with respect to the value predicted for λ_1_ (Supplementary Figure S7 B). The third eigenvalue of scaled PCA was close to the approximation provided by RMT (Supplementary Figure S7 D). To study cases in which the separation condition was not verified, we considered smaller number of genotypes (*L* ≤ 1000) and lower values of drift coefficients (*F_k_* ≤ 10%). For small values of *n* and *L*, a significant proportion of simulated data sets did not verify the separation condition (Supplementary Figure S8), although models were correctly specified. Those results provided additional evidence of biases in analyses of population structure with small data sets. Extending our simulation study to larger number of populations, we checked that the accuracy of the approximation of 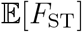 by the sum of the first (*K* – 1) PCA eigenvalues was proportional to *K*/*L* when the separation condition was verified (Supplementary Figure S9). Poorer accuracy was observed when the data failed to verify the separation condition.

### Human data

To provide evidence that the relationships between PCA eigenvalues and *F*_ST_ are verified for real genotypes, we computed *F*_ST_, their approximation by PCA, and the leading eigenvalues of the residual matrix for pairs and triplets of human population samples from The 1000 Genomes Project [50] (Table 1 and Supplementary Table S1). In pairwise analyses excluding admixed samples, the separation condition was verified in all analyses at the exception of the pair CEU-IBS, formed of two closely related European samples. The leading eigenvalue of scaled PCA was accurately approximated by 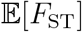, and the leading eigenvalue of the residual matrix was accurately predicted by RMT. For triplet analyses without admixed samples, the separation condition was also verified and the sum of the first two eigenvalues of scaled PCA was accurately approximated by 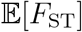. RMT still predicted the leading eigenvalue of the residual matrix accurately. In pair and triplet analyses including admixed samples, the separation condition was fulfilled in all analyses at the exception of the pair ACB-ASW (Table S1). The approximation of 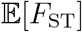 by the leading eigenvalues of scaled PCA was less accurate than in analyses without admixed samples. In the CEU-ASW analysis for example, 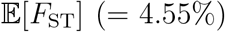 was lower than the leading eigenvalue of PCA (= 4.87%). An explanation for these discrepancies may be that *F*_ST_ is informative about the admixture proportion between admixed populations and their parental source populations [51]. With admixed samples, mismatches were also observed between the leading eigenvalue of the residual matrix and its prediction from RMT. The results suggest that the data do not agree with models of *K* discrete populations, and modified definitions of *F*_ST_ could be more appropriate for describing population structure in the presence of admixed individuals [52, 53].

**Table 1.** *F*_ST_ estimates for populations from The 1,000 Genomes Project.

| | Lead. eigen.<br>of PCA * | $F_{ST}$<br>across loci | Lead. eigen.<br>res. matrix** | RMT<br>approximation*** |
| --- | --- | --- | --- | --- |
| **CHB-CEU** | 5.65% | 5.65% | 0.42% | 0.48% |
| **CHB-YRI** | 8.35% | 8.35% | 0.36% | 0.37% |
| **CEU-YRI** | 7.21% | 7.21 % | 0.35% | 0.37% |
| **CEU-IBS** | 0.41% | 0.38 % | 0.37% | 0.41% |
| **CEU-YRI-CHB** | 9.99% | 9.98 % | 0.24% | 0.26% |
| **CEU-ASW** | 4.87% | 4.55 % | 0.75% | 0.52% |
| **CEU-YRI-ASW** | 6.12% | 5.82 % | 0.44% | 0.29% |

\* Sum of the leading eigenvalues of the PCA
\*\* Sum of the leading eigenvalues of the within-population (residual) matrix
\*\*\* RMT approximation for the leading eigenvalue of the within-population matrix
**IBS**: Iberian ( $n = 147$ ), **CHB**: Han Chinese in Beijing ( $n = 100$ ), **YRI**: Yoruba ( $n = 158$ ), **CEU**: Utah residents with European ancestry ( $n = 104$ ). **ASW**: Americans of African Ancestry in SW USA ( $n = 97$ ).

### Ancient DNA data

We illustrated how spectral estimates can be used to evaluate inbreeding coefficients from genotypes obtained after correction for experimental effects. We studied ancient DNA samples from early farmers from Anatolia (EFA, *n* = 23), steppe pastoralists from the Yamnaya culture (Steppe, *n* = 15), Western hunter-gatherers from Serbia (WHG, *n* = 31), and included Bell-Beaker samples from England and Germany (BKK, *n* = 38) [17, 54, 55, 56]. To estimate *F*_ST_ from those samples, we performed adjustment of pseudo-haploid genotypes for genomic coverage and for temporal distortions created by genetic drift. Temporal distortions were not expected to modify the average value of *F*_ST_ across loci. After genotypes were adjusted for coverage and corrected for distortions due to differences in sample ages, the resulting values could no longer be interpreted as allelic frequencies. The adjusted estimates for *F*_ST_ were equal to 4.7% for the *EFA – Steppe* data set, 5.8% for *EFA – WHG*, 5.1% for *Steppe – WHG* (Table 2). The separation condition was verified in all comparisons, and there was evidence of population structure in all pairwise analyses. Although individual PCA scores were impacted by coverage and temporal distortions (Figure 5), those unwanted effects did not generate substantial bias for PCA eigenvalues, leaving us with *F*_ST_ estimates that were similar with or without adjustment. For a three-population model including the EFA, WHG, and Steppe samples, the adjusted estimate for *F*_ST_ was equal to 7.0%, slightly lower than the uncorrected estimate (7.2 %, Table 2) and than the sum of the leading values of PCA (equal to 7.3%). The smallest eigenvalue of the between-population matrix was larger (2.6%) than the leading eigenvalue of the residual matrix (1.8%). This was no longer the case when the Bell Beakers from England and Germany were included in the data set. With Bell Beaker samples, the smallest eigenvalue of the between-population matrix was lower (1.0%) than the leading eigenvalue of the residual matrix (1.0%, Table 2). An explanation for this result is that the shared ancestry of Bell Beaker individuals [56] made PCA results inconsistent with a four-population model.

**Table 2.** *F*_ST_ estimates for ancient Eurasian samples with correction for genomic coverage.

| | $F_{ST}$ without<br>correction | $F_{ST}$ with<br>correction | Lead. eigen.<br>res. matrix* | RMT<br>threshold** |
| --- | --- | --- | --- | --- |
| **EFA-Steppe** | 4.8% | 4.7% | 3.1% | 2.8% |
| **EFA-WHG** | 5.9% | 5.8% | 3.3% | 2.0% |
| **Steppe-WHG** | 5.2% | 5.1 % | 3.8% | 2.3% |
| **EFA-Steppe-WHG** | 7.2% | 7.0% | 1.8 % (2.6%) | 1.5 % |
| **EFA-Steppe-WHG-BBK** | 5.9 % | 5.8% | 1.8 % (1.0%) | 1.0 % |

**EFA:** Early Farmers from Anatolia, **WHG:** Western Hunter-Gatherers, **Steppe:** Yamnaya pastoralists, **BBK:** Bell Beakers from England and Germany.
\* Leading eigenvalue of the within-population residual matrix (smallest eigenvalue of the between-population matrix)
\*\* RMT threshold for evidence of population structure in pairs: $(1/\sqrt{L} + 1/\sqrt{n-1})^2$ , RMT approximation in triplet and quadruplet: $(1 - \mathbb{E}[F_{ST}])(1/\sqrt{L} + 1/\sqrt{n-K})^2$ . $L$ : number of loci, $n$ : sample size.

**Figure 5.**
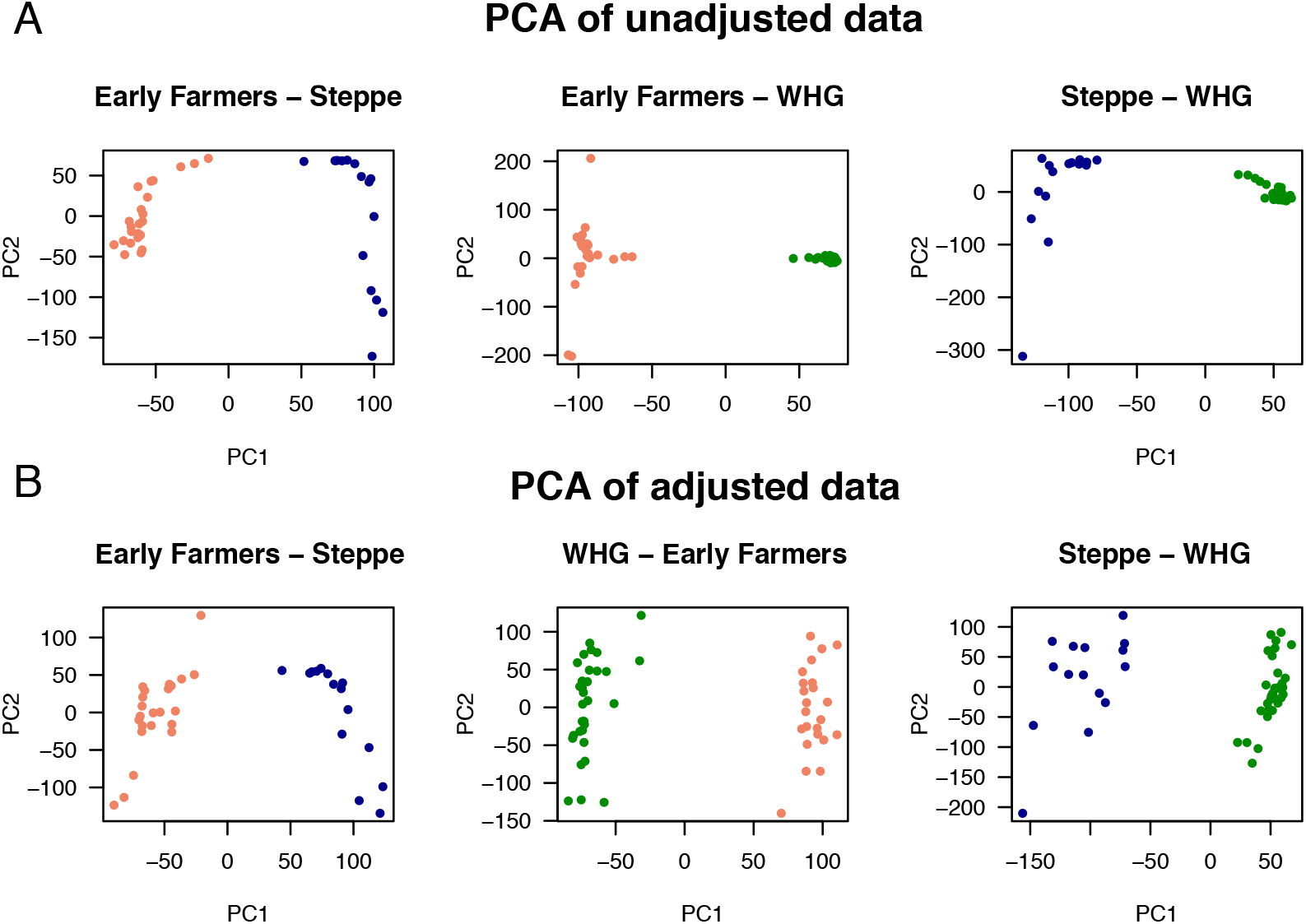
Correction for coverage in PC plots for pairs of ancient population samples. **(A)** PCA of unadjusted genotypes. **(B)** PCA of non-binary genotypic data adjusted for coverage. Population samples: Early Farmers (salmon color, *n*_1_ = 23), Steppe pastoralists (blue color, *n*_2_ = 15), (Western hunter gatherers, green color, *n*_3_ = 31)

### Genetic differentiation explained by bioclimatic factors

To provide a second illustration of the use of spectral estimates of inbreeding coefficients, we studied the role of bioclimatic factors in shaping population genetic structure in plants [29]. Here, the objective was to evaluate the proportion of differentiation explained by temperature and precipitation, which may influence adaptive genetic variation in those taxa. For 241 Swedish accessions of *Arabidopsis thaliana* taken from The 1,001 Genomes database [30], population structure was first evaluated by using a spatially explicit Bayesian algorithm. The individuals were clustered in two groups located in southern and northern Sweden (Figure 6A). For the groups estimated by spatial population structure analysis, the mean value of *F*_ST_ across loci was equal to 7.9%. This value was larger than the leading eigenvalue of the within-population matrix, equal to 4.9 %. The proportion of variance explained by the first PCA axis was equal to 8.5%, greater than *F*_ST_ (Figure 6). An explanation for this discrepancy is that a two-population model may not fit the data accurately, as PCA axes can capture spatial genetic variation unseen by the discrete population model. Population structure was further evaluated by using *K* = 3 ancestral populations. Southern individuals were split into two groups along a East-West axis, exhibiting mixed ancestry from those groups (Supplementary Figure S10). With three groups, the leading eigenvalues of the between-population matrix were equal to 7.8 % and 2.5%. The second eigenvalue was lower than the leading eigenvalue of the within-population matrix, equal to 3.7 %. According to those results, we decided to focus on information carried by a two-population model.

**Figure 6.**
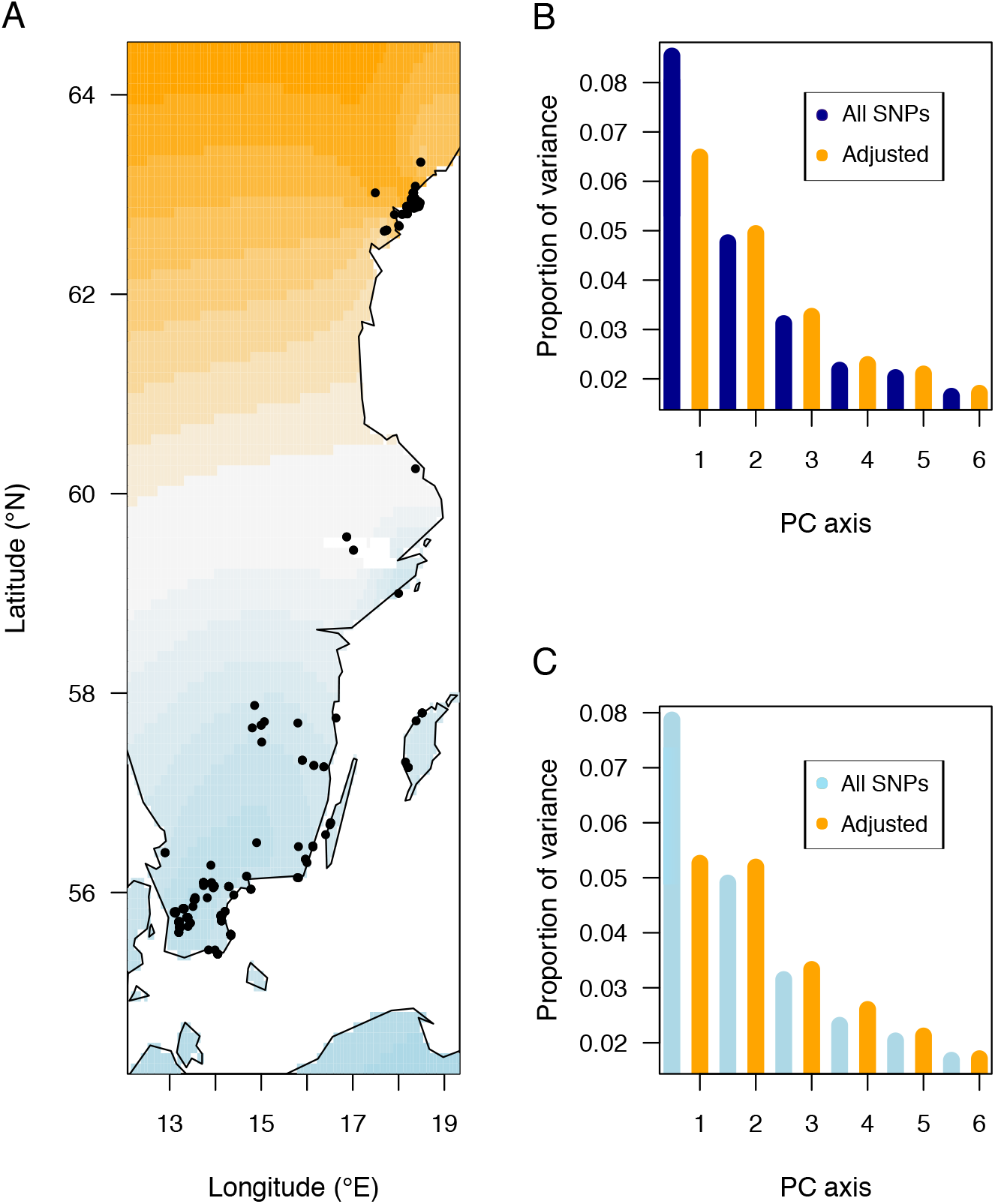
Neutral *F*_ST_ for *Arabidopsis thaliana* in Scandinavia. **(A)** Geographic locations of 241 samples and inference of population structure from a spatial method (Blue color: Southern cluster, Orange color: Northern cluster). **(B)** Proportion of variance explained by PC axes before adjustment of genotypes for bioclimatic variables (blue color) and after adjustment (orange color). **(C)** Proportion of variance explained by the first axis of the between-population matrix, and by the first axes of the residual matrix (five components) before adjustment for bioclimatic variables (blue color) and after adjustment (orange color). Wright’s coefficients are represented by the values for the first axis.

After adjusting for bioclimatic variation, the leading eigenvalue of the PCA was equal to 6.5% (Figure 6C). The eigenvalue of the between-population matrix – which defines the average value of *F*_ST_ for the adjusted genotypes, was equal to 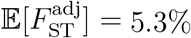. The second and subsequent PCA eigenvalues were equal to 4.9%, 3.2%, 2.3%, and those values were unaffected by the bioclimatic variables (Figure 6B). In addition, those eigenvalues agreed with the largest eigenvalues of the residual matrix, 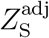, which were equal to 5.1%, 3.3%, 2.6% (Figure 6B). Comparing 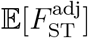 to 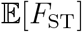 [42], these results showed that the relative proportion of variation explained by climate along the first axis was around 33%. The results provide evidence that climate had an impact on the differentiation of populations along the south-north axis, but had less influence on other axes of genetic variation. In summary, those numbers suggest that bioclimatic conditions played a major role in driving genetic divergence between northern and southern populations of Scandinavian *A. thaliana*.

### Conclusions

Assuming a model with *K* discrete populations, our study established a relationship between Wright’s inbreeding coefficient, *F*_ST_, and the (*K* – 1) leading eigenvalues of the between-population matrix and of scaled PCA. A similar relationship was established between Nei’s among-population diversity, *D*_ST_, and the leading eigenvalues of unscaled PCA. Those relationships justify the use of PCA to describe population genetic structure from large genotype matrices. They extend results obtained from coalescent theory for two divergent populations in Ref. [16] to any discrete population model. Assuming that RMT holds for residual matrices, they increase the accuracy of previous results, clarifying for which sample sizes and number of loci they could be valid. Computing the eigenvalues of the between-population matrix and of the residual matrix can be done numerically with a computing cost similar to PCA (Figure 1). In our simulations, we found that the leading eigenvalue of the residual matrix was well predicted by RMT. RMT also provided a threshold value of *F*_ST_, equal to 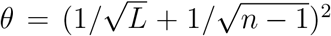 below which there is no evidence of population structure for two or more populations. This threshold differs from the threshold value of 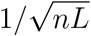 proposed in Ref. [10], and it was supported by simulations of single population models. In addition to connecting the PCA of a genotype matrix to inbreeding coefficients and related quantities, our results have several implications for the analysis of adjusted genotypes, providing statistics analogous to *F*_ST_ and *Q*_ST_ for those data. Adjusted genotypes arise in many applications, such as ancient DNA, to correct for biases due to technical or sampling artifacts, or in ecological genomics where it allows evaluating the part of population differentiation explained by environmental variation. The proposed estimates of inbreeding coefficients are thus of great importance to the understanding of the demographic history of populations and their adaptation to environmental variation.

## Methods

### PCA and SVD

For a genotype matrix **X** with *L* loci, centered PCA computes the eigenvalues, 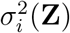, of the empirical covariance matrix, **ZZ**^*T*^/*n*, where **Z** = **Z**^c^ is the centered matrix, for which the mean value of each column has been substracted from **X** [9, 26]. Scaled PCA computes the eigenvalues, 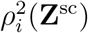, of the empirical correlation matrix, **Z**^sc^(**Z**^sc^)^*T*^/*n*, obtained for **Z**^sc^, the matrix in which each column of **Z** is divided by 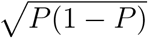. In order to obtain unbiased estimates, empirical covariance and correlation matrices are usually divided by (*n* – 1) instead of *n*. To avoid this complication, we kept *n* in all theoretical analyses (assuming that *n* is large). Unbiased estimates were used in empirical and simulated data analyses. Using the equivalence between PCA and SVD, the eigenvalues of PCA were computed as the squared non-null singular values of the matrix 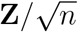.

### Spectral analysis

To make arguments easier to follow, we developed the analysis of eigenvalues for unscaled PCA. Extension to scaled PCA does not create mathematical complications but has heavier notations. This paragraph sketches the key arguments for the main result. More details are provided in *Supplementary Information*. We found that the squared Hilbert-Schmidt norm of the between-population matrix **Z**_ST_ is equal to

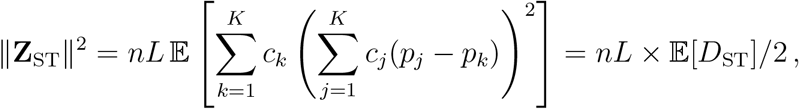

where the mathematical symbol 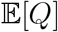 denotes the mean value of a quantitative or discrete quantity *Q* over the *L* loci. For scaled PCA, the squared norm is equal to 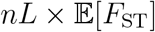. The matrices **Z**_ST_ and **Z**_S_ satisfy orthogonality conditions. In particular, the matrices are related by the Pythagorean equation ∥**Z**∥^2^ = ∥**Z**_ST_∥^2^ + ∥**Z**_S_∥^2^. This equation is the main argument for the alternative form of Theorem 1. In addition, stronger orthogonality conditions hold and enable showing that the accuracy of approximation of eigenvalues is of order 1/*L*.

### *F*-models

*F*-models are models for *K* discrete populations diverging from an ancestral gene pool [33]. In the ancestral gene pool, the derived allele is present with frequency *p*_anc_. The *K* populations diverged from each other and from the ancestral population with drift coefficient equal to *F_k_* relative to the ancestral pool. Conditional on *p*_anc_, the allele frequency at a particular locus in population *k* follows a beta distribution of shape parameters *p*_anc_(1 – *F_k_*)/*F_k_* and (1 – *p*_anc_)(1 – *F_k_*)/*F_k_*. To create a distribution over the *L* loci, *p*_anc_ is drawn from a beta distribution with shape parameters *a* and *b*, leading to 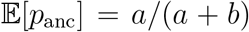. The expected ancestral heterozygozity, *H*_A_, is equal to 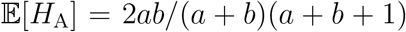. For *F*-models, the expected value of *D*_ST_ can be formulated as 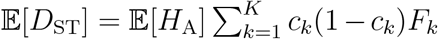 (*Supplementary Information*). Numerical values for 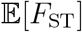 are less explicit, but they can be obtained by using Monte-Carlo simulations.

Simulations of *F*-models were performed in the R programming language. We performed simulations of single population models (*K* = 1) to check whether approximations derived from RMT appropriately describe the leading eigenvalue of scaled PCA in the absence of population structure. Simulations of *F*-models were performed with a value of the drift coefficient equal to *F* = 15%. The ancestral frequency for the derived allele, *p*_anc_, was drawn from a beta distribution with shape parameters *a* = 1 and *b* = 9, so that 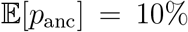 (Supplementary Figure S1). Simulations of *F*-models were performed with *K* = 2 to check whether the data could fit theoretical expectations for *D*_ST_ and *F*_ST_. Two hundred simulations of *F*-models were performed with equal values of the drift coefficients randomly drawn between 1% and 75% (*F*_1_ = *F*_2_). The ancestral frequency for the derived allele, *p*_anc_, was drawn from a beta distribution with shape parameters *a* = 1 and *b* = 4, so that 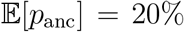. The total sample size was equal to *n* = 100 and the sample proportion *c*_1_ was drawn from a uniform distribution between 10% and 50%. Next, we considered three-population *F*-models with equal sample sizes and ancestral allele frequencies distributed according to the uniform distribution, (*a* = 1 and *b* = 1). With the uniform distribution, we found that 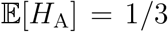, and the nonnull eigenvalues of the between-population covariance matrix could be computed as 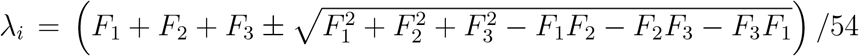, for *i* = 1, 2 (*Supplementary Information*). We had 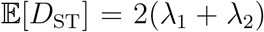. Two hundred simulations of three-population models were performed with unequal drift coefficients drawn between 1% and 25%. The total sample size was equal to *n* = 100 and the number of loci was equal to *L* = 20, 000. For values of *n* between 30 and 300, and number of loci between 100 and 1000, we performed additional simulations with small drift coefficients (*F_k_* ≤ 10%) to evaluate the probability that the data verify the separation condition. We also performed simulations of *F*-models for *K* between 4 and 40, using *n_k_* = 20 individuals in each sample (*L* = 20,000) and two models of drift: *F_k_* = 0.1 and *F_k_* = 0.2/*k* for all *k* = 4,…, 40.

### Approximation of the residual matrix from Random Matrix Theory (RMT)

For discrete population models, approximations of eigenvalues for the within-population (residual) matrix were obtained from RMT [10, 57, 58, 59, 60]. RMT considers large sample sizes, and keeps the ratio of the number of loci to the sample size at a constant value, *γ* = *L*/*n*. For single population models, we have **Z** = **Z**_S_, and the proportions of variance explained by each principal axis were approximated by the Marchenko-Pastur probability density function described by

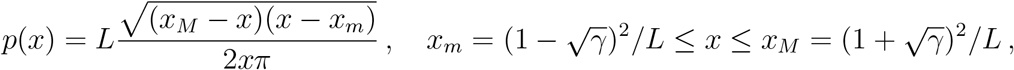

and the proportion of variance explained by the first principal axis was approximated by 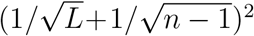. For *K* > 1, the Marchenko-Pastur density modelled the bulk distribution of eigenvalues for the within-population (residual) matrix. Under the separation condition (7), the proportion of variance explained by the *K*th principal axis was approximated by 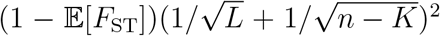. Regarding unscaled PCA, the largest singular value of the within-population matrix was approximated by 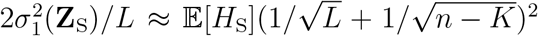. If there truly is a single population represented in the total sample, then *F*_ST_ for two equal size samples should be of order 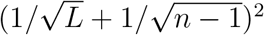.

### Human DNA analyses

We computed PCA eigenvalues, the mean values of *F*_ST_ and the leading eigenvalues of the residual matrix for pairs and triplets including human population samples from The 1000 Genomes Project [50]. In these comparisons, the number of SNPs was *L* ≈ 1.3M, after filtering out for minor allele frequency less than 5%. We considered samples from Han Chinese in Beijing (CHB, *n* = 100), Yoruba (YRI, *n* = 158), Utah residents with European ancestry (CEU, *n* = 104), Iberian (IBS, *n* = 147). We considered samples from populations with mixed ancestry, Americans of African Ancestry in SW USA (ASW, *n* = 97), Colombians from Medellin Colombia (CLM, *n* = 102), Puerto Ricans from Puerto Rico (PUR, *n* = 94), individuals of Mexican Ancestry from Los Angeles USA (MXL, *n* = 100), and African Caribbeans in Barbados (ACB, *n* = 98). Some pairs and triplets included individuals with mixed ancestry (while some other did not).

### Ancient DNA analyses

We analyzed 143,081 pseudo-haploid SNP genotypes from ancient samples of early farmers from Anatolia (*n* = 23), steppe pastoralists from the Yamnaya culture (*n* = 15), Western hunter-gatherers from Serbia (*n* = 31), and Bell-Beakers from England and Germany (*n* = 38). The data were extracted from a public data set available from David Reich lab’s repository (reich.hms.harvard.edu) [54, 17, 55, 56]. The ancient samples had a minimum coverage of 0.25x, a median coverage of 2.69x (mean of 2.98x) and a maximum coverage of 13.54x. Genotypes were adjusted for coverage by fitting a latent factor regression model with the number of factors equal to the number of sample minus two. The matrix was then adjusted for distortions due to differences in sample ages, resulting in surrogate genotypes encoded as continuous variables without any direct interpretation in terms of allelic frequency [28].

### Genomic and bioclimatic data analyses

We studied 241 swedish plant accessions from The 1,001 Genomes database for *Arabidopsis thaliana* [30]. A matrix of SNP genotypes was obtained by considering variants with minor allele frequency greater than 5% and a density of variants around one SNP every 1,000 bp (167,475 SNPs). The individuals were clustered in groups based on analysis of population structure accounting for geographic proximity [61]. Global climate and weather data corresponding to individual geographic coordinates were downloaded from the World-Clim database (https://worldclim.org). Eighteen bioclimatic variables, derived from the monthly temperature and rainfall values, were considered as representing the current environmental matrix. Correction of genotypes for locus-specific effects of the eighteen environmental variables was performed with a latent factor regression model implemented in the R package `lfmm` [41]. For the matrix of centered genotypes, **Z**, and the matrix of eighteen bioclimatic variables, **Y**, the program estimated a matrix of surrogate genotypes, **W**, by adjusting a regression model of the form **Z** = **YB**^*T*^ + **W** + *ϵ*. To keep the latent matrix estimate (**W**) as close as possible to **Z**, we used *k* = *n* – 19 = 222 factors to compute **W**.

## Data availability

The data used in our study were publicly available from their cited reference.

## Code availability

All codes necessary to reproduce the simulations and data analyses presented in our study are in an R package available from the following link: https://github.com/bcm-uga/spectralfst

---

**Box 1.** Notations

Box 1. Notations
|  |  |
| --- | --- |
| $n$ | Sample size |
| $L$ | Number of genomic loci |
| $F_{ST}$ | Wright's fixation index, computed from Nei's formula with correction for unequal sample sizes |
| $H_S$ | Within-population genetic diversity |
| $H_T$ | Genetic diversity in the total population |
| $D_{ST}$ | Among (or between) population genetic diversity |
| $\mathbf{X}$ | Matrix of SNP genotypes for $n$ individuals at $L$ loci |
| $\mathbf{P}$ | Vector of SNP frequency for the $L$ loci |
| $\mathbf{Z}$ | Matrix of centered genotypes, $\mathbf{X} - \mathbf{P}$ |
| $\mathbf{Z}^{sc}$ | Matrix of scaled genotypes, $\mathbf{Z}/\sqrt{\mathbf{P}(1 - \mathbf{P})}$ |
| $\mathbf{Z}_{ST}$ | An $n \times L$ matrix describing between-population data repeated for individuals from a same population |
| $\mathbf{Z}_S$ | An $n \times L$ matrix describing within-population data |
| $\sigma_k^2(\mathbf{Z})$ | Eigenvalues of the empirical covariance matrix (unscaled PCA) |
| $\rho_k^2(\mathbf{Z}^{sc})$ | Eigenvalues of the empirical correlation matrix (scaled PCA), also equal to $L$ times the proportions of variance explained by the principal axes |

## Notes

### Competing Interest Statement

The authors have declared no competing interest.

### Summary of Updates

Section on Theory updated to clarify hypotheses, Supplemental files updated, new examples added

