## supplementary information for "A Spectral Theory for Wright’s Inbreeding Coefficients and Related Quantities"

Olivier François Clément Gain

Université Grenoble-Alpes, Centre National de la Recherche Scientifique, Grenoble INP, TIMC-IMAG CNRS UMR 5525, 38000 Grenoble, France.

**4 Supplementary Tables and Figures** **14**

### 1 Inbreeding coefficients and matrix norms

#### 1.1 Results for $D_{\mathrm{ST}}$ and $F_{\mathrm{ST}}$

Consider $K$ discrete populations and a sample of $n$ organisms from those populations, where $n_k$ individuals originate from population $k$. Obviously, we have $n_1 + \cdots + n_K = n$. The coefficient $c_k = n_k/n$ represents the proportion of individuals sampled from population $k$. Genetic variation is observed at a diallelic locus having a reference allele and a derived allele. The frequency of the derived allele in population $k$ is equal to $p_k$. The frequency of the reference allele in population $k$ is equal to $q_k = 1 - p_k$. The frequency of the derived allele in the total sample is then equal to $P = \sum_{k=1}^{K} c_k p_k$. The total heterozygozity is defined as $H_{\mathrm{T}} = 2P(1-P)$, and we set $H_{\mathrm{S}} = 2\sum_{k=1}^{K} c_k p_k q_k$. Using notations introduced in [8, 5, 6], $D_{\mathrm{ST}}$ is defined as the difference $D_{\mathrm{ST}} = H_{\mathrm{T}} - H_{\mathrm{S}}$, and is equal to

$$D_{\mathrm{ST}} = 2\left(\sum_{j=1}^{K}\sum_{k=1}^{K} c_j c_k p_j q_k - \sum_{j=1}^{K} c_j p_j q_j\right) .$$

Rearranging terms on the left-hand side of the above equation, we find that

$$D_{\mathrm{ST}} = 2\left(\sum_{j=1}^{K} c_j p_j \sum_{k=1}^{K} c_k (p_j - p_k)\right) = 2\sum_{j<k} c_j c_k (p_j - p_k)^2 .$$

Now, developing the expression

$$A = \sum_{k=1}^{K} c_k \left(\sum_{j=1}^{K} c_j (p_j - p_k)\right)^2 ,$$

we find that $D_{\mathrm{ST}}$ is also equal to

$$D_{\mathrm{ST}} = \sum_{k=1}^{K} c_k \left(\sum_{j=1}^{K} c_j (p_j - p_k)\right)^2 .$$

#### 1.2 Details on matrix norms

For any matrix, $\mathbf{Y}$, of dimension $n \times L$, the Hilbert-Schmidt norm – or Frobenius norm – of $\mathbf{Y}$ is defined by

$$\|\mathbf{Y}\| = \left( \sum_{i=1}^{n} \sum_{\ell=1}^{L} y_{i\ell}^2 \right)^{1/2} .$$

In particular, the squared norm of $\mathbf{Y}$ corresponds to the sum of the eigenvalues of $\mathbf{Y}^T\mathbf{Y}$

$$\|\mathbf{Y}\|^2 = \mathbf{Tr}(\mathbf{Y}^T\mathbf{Y}) = n \sum_{i=1}^{\min(n,L)} \sigma_i^2(\mathbf{Y}) \, ,$$

where $\sigma_i(\mathbf{Y})$ is a non-null singular value of $\mathbf{Y}/\sqrt{n}$, and $\mathbf{Tr}(\mathbf{M})$ denotes the trace of the matrix $\mathbf{M}$. For all $i$, $\sigma_i^2(\mathbf{Y})$ is a non-null eigenvalue of $\mathbf{Y}\mathbf{Y}^T/n$ and of $\mathbf{Y}^T\mathbf{Y}/n$. We shall use the term *eigenvalue* as a synonymous of *squared singular value.*

### 2 Main results

#### 2.1 Proof of Theorem 1

Let $\mathbf{X} = (x_{i\ell})$ record genotypes at $L$ loci for $n$ individuals sampled from $K$ discrete populations. For haploids, we have $x_{i\ell} = 0, 1$, and for diploids, we have $x_{i\ell} = 0, 1, 2$. We consider autosomes, and we suppose that a diploid genome is formed of two haploid genomes. When the phase of a diploid genotype is unknown, we use an arbitrary phase, either defined as $(0, 1)$ or as $(1, 0)$, in order to create two haploid genotypes. With this simplification, the diploid case amounts to consider a sample of $2n$ haploid individuals. Let $\mathbf{Z}^{\mathrm{c}} = \mathbf{X} - \mathbf{P}$ be the centered matrix, for which the mean value of a column is null. Let $\mathbf{Z}^{\mathrm{sc}}$ be the scaled matrix in which each column has variance is equal to one, that is, $\mathbf{z}_{i\ell}^{\mathrm{sc}} = \mathbf{z}_{i\ell}^{\mathrm{c}}/\sqrt{P_\ell(1 - P_\ell)}$ [7]. For a clearer presentation, we only describe results for $\mathbf{Z}^{\mathrm{c}}$ and omit the superscript. The same principles and results also apply to $\mathbf{Z}^{\mathrm{sc}}$ in a straightforward way.

The $n \times L$ matrix $\mathbf{Z}$ can be decomposed as the sum of between and within-population matrices, $\mathbf{Z} = \mathbf{Z}_{\mathrm{ST}} + \mathbf{Z}_{\mathrm{S}}$, as follows. For an individual $i$ sampled from population $k$ and an arbitrary locus $\ell$, the between-population matrix $\mathbf{Z}_{\mathrm{ST}}$ has general term

$$z_{i\ell}^{st} = \sum_{j=1}^{K} c_j (p_{k\ell} - p_{j\ell}) \,,$$

where this term is repeated $n_k$ times at each locus, for all $i$ in population $k$. The allele frequencies, $p_{k\ell}$, are specific to each locus. The within-population matrix $\mathbf{Z}_{\mathrm{S}}$ has general term

$$z_{i\ell}^{s} = x_{i\ell} - p_{k\ell} \,.$$

Here we compute the matrix norms of the between and within-population matrices separately. For the between-population matrix, we have

$$\|\mathbf{Z}_{\mathrm{ST}}\|^2 = \sum_{i=1}^{n} \sum_{\ell=1}^{L} (z_{i\ell}^{st})^2 \,,$$

which is equal to

$$\|\mathbf{Z}_{\mathrm{ST}}\|^2 = nL\,\mathbb{E}\left[\sum_{k=1}^{K} c_k \left(\sum_{j=1}^{K} c_j (p_j - p_k)\right)^2\right] = nL\,\mathbb{E}[D_{\mathrm{ST}}]/2 \,.$$

We use the notation $\mathbb{E}[Q] = \sum_{\ell=1}^{L} Q_\ell / L$ everywhere to denote the average value of $Q_\ell$ over all loci. For the within-population matrix, we have

$$\|\mathbf{Z}_{\mathrm{S}}\|^2 = \sum_{i=1}^{n} \sum_{\ell=1}^{L} (z_{i\ell}^{s})^2 \,,$$

which is equal to

$$\|\mathbf{Z}_{\mathrm{S}}\|^2 = nL\,\mathbb{E}\left[\sum_{k=1}^{K} c_k \mathrm{Var}(x_i|i \in \mathrm{pop}_k)\right] = nL\,\mathbb{E}[H_{\mathrm{S}}]/2\,.$$

The scaled matrix, $\mathbf{Z}^{\mathrm{sc}}$, can be obtained from $\mathbf{Z}$ by dividing each column by $\sqrt{P_\ell(1-P_\ell)}$. Thus the squared norm of $\mathbf{Z}^{\mathrm{sc}}_{\mathrm{ST}}$ is equal to

$$\|\mathbf{Z}^{\mathrm{sc}}_{\mathrm{ST}}\|^2 = nL\,\mathbb{E}[F_{\mathrm{ST}}]\,.$$

This important result redefines the average value $\mathbb{E}[F_{\mathrm{ST}}]$ from the spectrum of the between-population matrix. The definition is valid regardless of sample size, number of loci and separation of between and within-components.

#### 2.2 Proof of Corollary

Consider a matrix of $L$ haploid genotypes for $n$ individuals, $\mathbf{X}$. Centered PCA computes the eigenvalues and eigenvectors of the empirical covariance matrix, $\mathbf{Z}\mathbf{Z}^T/n$. (In data analyses, we use $(n-1)$ instead of $n$, in order to manipulate unbiased estimates of covariance matrices. Since $n \approx n-1$ for large sample sizes, taking $n$ introduces a small bias but enable a lighter presentation.) The eigenvalues of the empirical covariance matrix correspond to the squared non-null singular values of $\mathbf{Z}/\sqrt{n}$ [4]. As in the above section, the matrix $\mathbf{Z}$ is decomposed into a sum of between and within-population components, $\mathbf{Z} = \mathbf{Z}_{\mathrm{ST}} + \mathbf{Z}_{\mathrm{S}}$. The matrix $\mathbf{Z}_{\mathrm{ST}}$ has $(K-1)$ non-null singular values. First, we note that the matrices $\mathbf{Z}_{\mathrm{ST}}$ and $\mathbf{Z}_{\mathrm{S}}$ satisfy the following orthogonality condition,

$$\mathbf{Z}^T_{\mathrm{ST}}\mathbf{Z}_{\mathrm{S}} = \mathbf{0}\,.$$

The orthogonality condition can be checked directly, by computing the product matrix coefficients for all $\ell, m$ in $1, \ldots, L$. Those coefficients are equal to

$$\sum_{i=1}^{n} z_{i\ell}^{st} z_{im}^{s} = n \sum_{k=1}^{K} c_k \left( \sum_{j=1}^{K} c_j (p_{k\ell} - p_{j\ell}) \right) \sum_{i \in \mathrm{pop_k}} (x_{im} - p_{km}) \,.$$

Remarking that we have

$$\sum_{i \in \mathrm{pop_k}} (x_{im} - p_{km}) = 0 \,,$$

for each $k$, the orthogonality condition is obviously verified.

Using the definition of the squared norm as the trace of a matrix, we now have a Pythagorean relationship for between and within-population matrices

$$\|\mathbf{Z}_{\mathrm{ST}} + \mathbf{Z}_{\mathrm{S}}\|^2 = \mathbf{Tr}(\mathbf{Z}_{\mathrm{ST}}^T \mathbf{Z}_{\mathrm{ST}}) + \mathbf{Tr}(\mathbf{Z}_{\mathrm{S}}^T \mathbf{Z}_{\mathrm{S}}) = \|\mathbf{Z}_{\mathrm{ST}}\|^2 + \|\mathbf{Z}_{\mathrm{S}}\|^2 \,.$$

This result concludes the proof of the corollary.

#### 2.3 Approximation of $D_{\mathrm{ST}}$ and $F_{\mathrm{ST}}$ by PCA eigenvalues

In this section, we provide arguments for the following result. Assume that 1) the smallest non-null singular value of $\mathbf{Z}_{\mathrm{ST}}$ is larger than the leading eigenvalue of $\mathbf{Z}_{\mathrm{S}}$, 2) following RMT, the leading eigenvalue of $\mathbf{Z}_{\mathrm{S}}/\sqrt{n}$ is of order $(1/\sqrt{n} + 1/\sqrt{L})^2$. Then, for any non-null singular value, $\sigma_k(\mathbf{Z}_{\mathrm{ST}})$, of $\mathbf{Z}_{\mathrm{ST}}/\sqrt{n}$, there exists a singular value, $\sigma_k(\mathbf{Z})$, of $\mathbf{Z}/\sqrt{n}$ such that we have

$$|\sigma_k^2(\mathbf{Z})/L - \sigma_k^2(\mathbf{Z}_{\mathrm{ST}})/L| \leq C/L \,, \quad k = 1, \ldots, K-1 \,.$$

For $\sigma = \sigma(\mathbf{Z}_{\mathrm{ST}})$, a non-null singular value of $\mathbf{Z}_{\mathrm{ST}}/\sqrt{n}$, and $\tilde{\sigma} = \tilde{\sigma}(\mathbf{Z}_{\mathrm{S}})$, a non-null singular value of $\mathbf{Z}_{\mathrm{S}}/\sqrt{n}$, the left singular vectors, $\mathbf{u}$ and $\tilde{\mathbf{u}}$, associated with $\sigma$ and $\tilde{\sigma}$ are orthogonal vectors. To see it, compute

$$n^2 \sigma^2 \tilde{\sigma}^2 \mathbf{u}^T \tilde{\mathbf{u}} = \mathbf{u}^T \mathbf{Z}_{\mathrm{ST}} (\mathbf{Z}_{\mathrm{ST}}^T \mathbf{Z}_{\mathrm{S}}) \mathbf{Z}_{\mathrm{S}}^T \tilde{\mathbf{u}}.$$

Since $\mathbf{Z}_{\mathrm{ST}}^T\mathbf{Z}_{\mathrm{S}} = \mathbf{0}$, the middle term is null and the left singular vectors, $\mathbf{u}$ and $\tilde{\mathbf{u}}$, satisfy $\mathbf{u}^T\tilde{\mathbf{u}} = 0$.

Assuming that $L/n = \gamma$ for large values of $L$ and $n$, we now show that a non-null singular value of the between-population matrix, $\mathbf{Z}_{\mathrm{ST}}/\sqrt{n}$, is an approximate singular value of the genotype matrix, $\mathbf{Z}/\sqrt{n}$. A similar result holds for the eigenvalues of the residual matrix, $\mathbf{Z}_{\mathrm{S}}/\sqrt{n}$. To show the result for the between-population matrix, let $\mathbf{u}$ be a left singular vector of $\mathbf{Z}_{\mathrm{ST}}/\sqrt{n}$ and let $\sigma$ be the corresponding non-null singular value. We have

$$\mathbf{Z}\mathbf{Z}^T\mathbf{u}/n = \mathbf{Z}_{\mathrm{ST}}\mathbf{Z}_{\mathrm{ST}}^T\mathbf{u}/n + \mathbf{Z}_{\mathrm{S}}\mathbf{Z}_{\mathrm{ST}}^T\mathbf{u}/n.$$

For the first term at the right-hand side, we have $\mathbf{Z}_{\mathrm{ST}}\mathbf{Z}_{\mathrm{ST}}^T\mathbf{u}/n = \sigma^2\mathbf{u}$. For the second term, we have $\mathbf{Z}_{\mathrm{ST}}^T\mathbf{u}/\sqrt{n} = \sigma\mathbf{v}$, where $\mathbf{v}$ is a right singular vector of $\mathbf{Z}_{\mathrm{ST}}^T/\sqrt{n}$, and

$$\mathbf{Z}\mathbf{Z}^T\mathbf{u}/n = \sigma^2\mathbf{u} + \sigma\mathbf{Z}_{\mathrm{S}}\mathbf{v}/\sqrt{n}\,.$$

By definition of the spectral radius, we have

$$\|\mathbf{Z}_{\mathrm{S}}\mathbf{v}/\sqrt{n}\|^2 \leq \mathrm{radius}(\mathbf{Z}_{\mathrm{S}}\mathbf{Z}_{\mathrm{S}}^T/n)\,.$$

Now, note that the eigenvalue $\sigma^2$ is of order $O(L)$ (we have $\sigma^2 < \|\mathbf{Z}_{\mathrm{ST}}\|^2/n = O(L)$ by Theorem 1). According to assumption 2) for $\mathbf{Z}_{\mathrm{S}}$, we have

$$\sigma^2\|\mathbf{Z}_{\mathrm{S}}\mathbf{v}/\sqrt{n}\|^2 = (1+\sqrt{\gamma})^2 \times O\left(1\right)\,.$$

Put together, the last equations imply that

$$\mathbf{Z}\mathbf{Z}^T\mathbf{u}/n = \sigma^2\mathbf{u} + O(1)\,.$$

Thus, for large $n$ and $L$, $\mathbf{Z}\mathbf{Z}^T\mathbf{u}/n \approx \sigma^2\mathbf{u}$ plus a negligible term, meaning that $\mathbf{u}$ is an approximate eigenvector of $\mathbf{Z}/\sqrt{n}$, and $\sigma^2$ is an approximate eigenvalue of

$\mathbf{Z}/\sqrt{n}$. Now, we consider the vector space spanned by the first $K-1$ eigenvectors of $\mathbf{C}_{\text{ST}} = \mathbf{Z}_{\text{ST}}\mathbf{Z}_{\text{ST}}^T/n$. Then, consider

$$\mathbf{C}_{\text{ST}}\mathbf{U} = \mathbf{U}\Sigma\,,$$

where $\Sigma = \text{diag}(\sigma_1^2, \ldots, \sigma_{K-1}^2)$ and $\mathbf{U} = (\mathbf{u}_1, \ldots, \mathbf{u}_{K-1})$ are the non-null eigenvalues and left singular vectors of $\mathbf{Z}_{\text{ST}}/\sqrt{n}$. For $\mathbf{C} = \mathbf{Z}\mathbf{Z}^T/n$, we have

$$\mathbf{U}^T\mathbf{C}\mathbf{U} = \Sigma + \mathbf{U}^T O(1)\,.$$

The condition number of $\mathbf{U}$ being equal to one, we can apply the Bauer-Fike Theorem [3, 2]. For each non-null singular value of $\mathbf{Z}_{\text{ST}}/\sqrt{n}$, $\sigma^2(\mathbf{Z}_{\text{ST}})$, there is a singular value of $\mathbf{Z}/\sqrt{n}$, $\sigma^2(\mathbf{Z})$, such that

$$|\sigma^2(\mathbf{Z}) - \sigma^2(\mathbf{Z}_{\text{ST}})| = O(1)\,.$$

Using the results on matrix norms established in the previous section, we have, on one hand

$$\|\mathbf{Z}_{\text{ST}}\|^2 = nL\mathbb{E}[D_{\text{ST}}/2],$$

while, on another hand, we have

$$\|\mathbf{Z}_{\text{ST}}\|^2 = \mathbf{Tr}(\mathbf{Z}_{\text{ST}}\mathbf{Z}_{\text{ST}}^T) = n\sum_{k=1}^{K-1}\sigma_k^2(\mathbf{Z}_{\text{ST}})\,.$$

Under the separation condition and the RMT hypothesis, we have

$$\sum_{k=1}^{K-1}\sigma_k^2(\mathbf{Z}) \approx \sum_{k=1}^{K-1}\sigma_k^2(\mathbf{Z}_{\text{ST}})\,,$$

and

$$L \times \mathbb{E}[D_{\text{ST}}]/2 = \sum_{k=1}^{K-1}\sigma_k^2(\mathbf{Z}_{\text{ST}}) \approx \sum_{k=1}^{K-1}\sigma_k^2(\mathbf{Z})\,.$$

For centered PCA, we obtained that $L\mathbb{E}[D_{\mathrm{ST}}]/2$ is close to the sum of the $(K-1)$ leading eigenvalues of $\mathbf{Z}/\sqrt{n}$. Turning to scaled PCA creates no complication. Since the empirical correlation matrix is obtained by scaling the columns of $\mathbf{Z}$, a spectral decomposition holds for the scaled matrix as well. For scaled PCA, the sum of the $(K-1)$ leading eigenvalues is equal to

$$L \times \mathbb{E}[F_{\mathrm{ST}}] = \sum_{k=1}^{K-1} \rho_k^2(\mathbf{Z}_{\mathrm{ST}}) \approx \sum_{k=1}^{K-1} \rho_k^2(\mathbf{Z})\,.$$

In other words, $\mathbb{E}[F_{\mathrm{ST}}]$ can be approximated by the proportion of variance explained by the first $(K-1)$ principal components of the genotype matrix. The accuracy of the approximation is expected to be of order $O(K/L)$ with respect to $K$ and $L$. This error term is supported by numerical simulations of $F$-models (Figure SI A, Figure S5 and Figure S9).

### 3 Mathematical analysis of $F$-models

#### 3.1 $F$-models

$F$-models are discrete population models in which $K$ populations diverged from an ancestral gene pool [1]. The models are without mutation, migration or selection. In the ancestral gene pool, the frequency of the derived allele is equal to $p_{\mathrm{anc}}$. The populations diverged from each other with population-specific drift coefficients, $F_k$, relative to the ancestral pool. Conditional on $p_{\mathrm{anc}}$, the allele frequency at a particular locus in population $k$ follows a beta distribution of shape parameters $p_{\mathrm{anc}}(1-F_k)/F_k$ and $(1-p_{\mathrm{anc}})(1-F_k)/F_k$. To create a distribution over the $L$ loci, $p_{\mathrm{anc}}$ is drawn from a beta probability density with shape parameters $a$ and $b$. The expected value for this distribution is equal to $\mathbb{E}[p_{\mathrm{anc}}] = a/(a+b)$. $F$-models parameters are chosen so that $\mathbb{E}[p_k|p_{\text{anc}}] = p_{\text{anc}}$ and $\text{Var}(p_k|p_{\text{anc}}) = p_{\text{anc}}(1-p_{\text{anc}})F_k$, for all $k$. For $F$-models, the expected ancestral heterozygozity is equal to

$$\mathbb{E}[H_{\text{A}}] = \int_0^1 2p_{\text{anc}}(1-p_{\text{anc}})f(p_{\text{anc}})dp_{\text{anc}} = \frac{2ab}{(a+b)(a+b+1)} \, .$$

#### 3.2 Expected values

In this section, we compute the expected values of $D_{\text{ST}}$, $H_{\text{S}}$ and $H_{\text{T}}$ under the $F$-model. Those results may not be new, but they are difficult to find from the existing literature. For the first quantity, we have $D_{\text{ST}} = 2\mathbb{E}[\sum_{j<k} c_j c_k (p_j - p_k)^2]$. Given $p_{\text{anc}}$, the conditional expectation of the squared allele frequency difference $(p_j - p_k)^2$ is equal to

$$\mathbb{E}[(p_j - p_k)^2|p_{\text{anc}}] = \mathbb{E}[(p_j - p_{\text{anc}})^2|p_{\text{anc}}] + \mathbb{E}[(p_k - p_{\text{anc}})^2|p_{\text{anc}}] \, .$$

The sum of squares arises from the interpretation of the $F$-model as a starlike tree model of divergence or more pragmatically from the conditional independence of the allele frequencies $p_j$ and $p_k$ given $p_{\text{anc}}$. This leads us to

$$\mathbb{E}[(p_j - p_k)^2|p_{\text{anc}}] = p_{\text{anc}}(1-p_{\text{anc}})(F_j + F_k) \, .$$

Thus we have

$$\mathbb{E}[\sum_{j \neq k} c_j c_k (p_j - p_k)^2|p_{\text{anc}}] = p_{\text{anc}}(1-p_{\text{anc}}) \sum_{j \neq k} c_j c_k (F_j + F_k) \, .$$

Reorganizing the sum, we find that

$$\sum_{j \neq k} c_j c_k F_j = \sum_{j=1}^{K} c_j F_j \sum_{k \neq j}^{K} c_k = \sum_{j=1}^{K} c_j (1 - c_j) F_j \, ,$$

and we can conclude that

$$\mathbb{E}[D_{\mathrm{ST}}] = \left( \sum_{k=1}^{K} c_k(1-c_k)F_k \right) \mathbb{E}[H_{\mathrm{A}}] \,.$$

Following similar calculations, the results for $H_{\mathrm{T}}$ and $H_{\mathrm{S}}$ are given by

$$\mathbb{E}[H_{\mathrm{T}}] = \left( 1 - \sum_{k=1}^{K} c_k^2 F_k \right) \mathbb{E}[H_{\mathrm{A}}] \,.$$

and

$$\mathbb{E}[H_{\mathrm{S}}] = \left( 1 - \sum_{k=1}^{K} c_k F_k \right) \mathbb{E}[H_{\mathrm{A}}] \,,$$

We did not obtain an explicit analytical formula for the expected value of $F_{\mathrm{ST}}$ under the $F$-model. For computing $\mathbb{E}[F_{\mathrm{ST}}]$, we relied on numerical integration performed by a Monte-Carlo method.

#### 3.3 Three-population models

For three-population $F$-models with equal sample sizes and for ancestral allele frequencies distributed according to the uniform distribution, beta($a = 1$, $b = 1$), the $3 \times 3$ matrix $\Lambda$ corresponding to the between-population covariance matrix for allele frequencies can be calculated as

$$\mathbf{\Lambda} = \frac{1}{162} \begin{pmatrix} 4F_1 + F_2 + F_3 & F_3 - 2(F_1 + F_2) & F_2 - 2(F_1 + F_3) \\ * & F_1 + 4F_2 + F_3 & F_1 - 2(F_2 + F_3) \\ * & * & F_1 + F_2 + 4F_3 \end{pmatrix} ,$$

where stars correspond to symmetric coefficients. After elementary linear algebra, the first eigenvalue of $\mathbf{\Lambda}$ can be computed as

$$\lambda_1 = \frac{1}{54} \left( F_1 + F_2 + F_3 + \sqrt{F_1^2 + F_2^2 + F_3^2 - F_1F_2 - F_2F_3 - F_3F_1} \right) .$$

The second eigenvalue is equal to

$$\lambda_2 = \frac{1}{54}\left(F_1 + F_2 + F_3 - \sqrt{F_1^2 + F_2^2 + F_3^2 - F_1F_2 - F_2F_3 - F_3F_1}\right) ,$$

and we have $\mathbf{Tr}(\mathbf{\Lambda}) = \lambda_1 + \lambda_2 = \mathbb{E}[D_{\mathrm{ST}}]/2$. At the cost of less readable formulas, these results could be extended to arbitrary values of $c_k$, $a$, $b$.

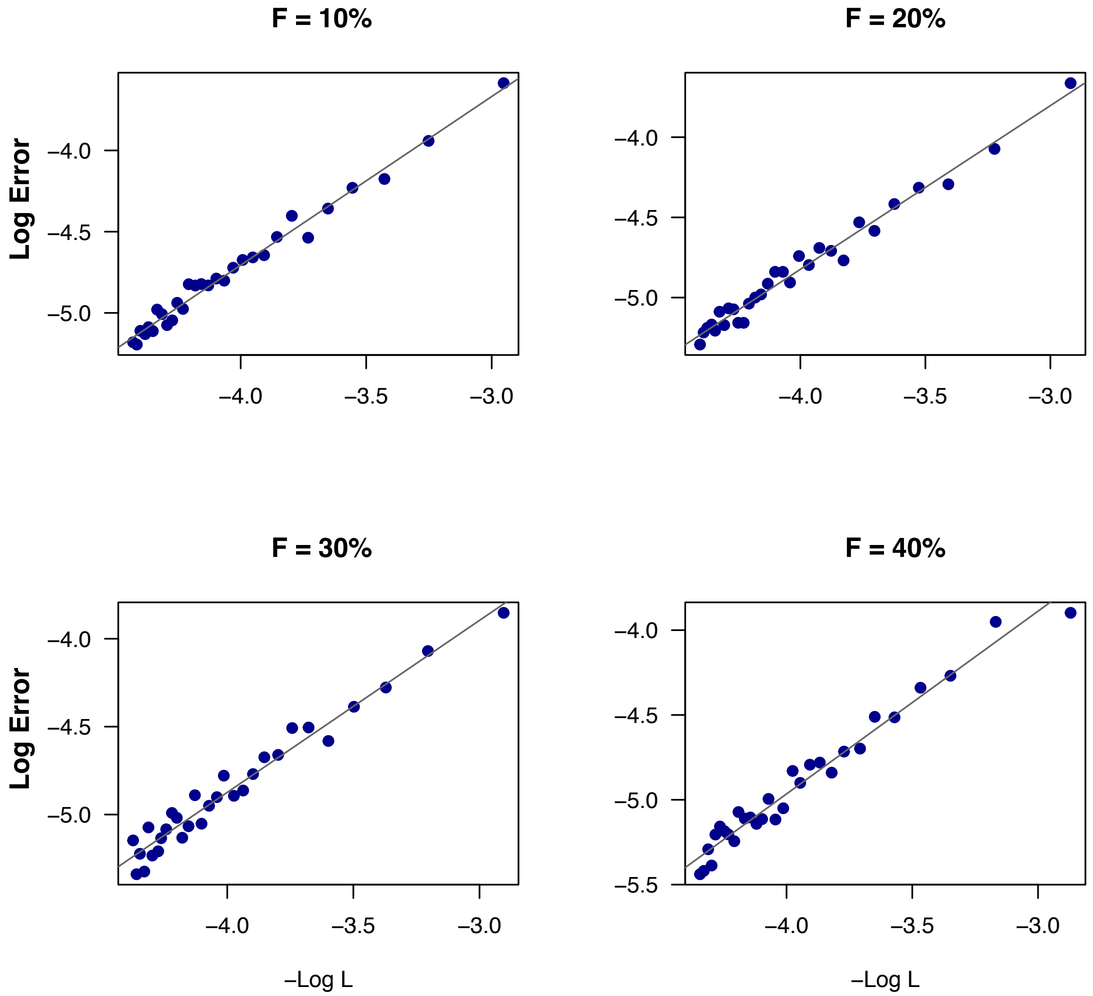

**Figure SI A. Order of approximation of $\mathbb{E}[F_{\mathrm{ST}}]$ by PCA leading eigenvalue in two-population models.** Difference between the leading eigenvalue and the average value of $F_{\mathrm{ST}}$ across loci as a function of the inverse of the number of loci $1/L$ on a log10-scale. Simulations of $F$-models were performed with $p_{\mathrm{anc}}$ drawn from a beta distribution with shape parameters $a = 1$ and $b = 4$, $n = 100$ individuals, and drift coefficients as indicated on the top of each panel.

### 4 Supplementary Tables and Figures

**Table S1.** $F_{\mathrm{ST}}$ estimates for populations from The 1,000 Genomes Project.

**Figure S1.** Random matrix theory approximation of proportions of variance in single population $F$-models.

**Figure S2.** Separation of variance components in artificial population samples built from single population $F$-models.

**Figure S3.** Accuracy of $D_{\mathrm{ST}}$ estimates with respect to their theoretical values in two-population models.

**Figure S4**. Accuracy of approximation of $F_{\mathrm{ST}}$ and separation condition in two-population $F$-models.

**Figure S5**. Approximation errors decrease as $1/L$ in two-population $F$-models.

**Figure S6.** First eigenvalue ($F_{\mathrm{ST}}$ estimate) as a function of sample size in two-population models.

**Figure S7.** Leading eigenvalues in three-population $F$-models.

**Figure S8.** Separation condition in three-population models.

**Figure S9**. Accuracy of approximation of $F_{\mathrm{ST}}$ by PCA for $K$-population $F$-models.

**Figure S10**. Estimates of ancestry coefficients for 241 Swedish accessions of *A. thaliana*.

**Table S1.** $F_{\mathrm{ST}}$ **estimates for populations from The 1,000 Genomes Project**

| | Lead. eigen. of PCA* | $F_{\mathrm{ST}}$ across loci | Lead. eigen. res. matrix** | RMT approximation*** |
|---|---|---|---|---|
| **YRI-IBS** | 7.27% | 7.27 % | 0.31% | 0.32% |
| **YRI-IBS-CHB** | 9.75% | 9.74 % | 0.25% | 0.25% |
| **ACB-ASW** | 1.26% | 0.60 % | 1.05% | 0.56% |
| **PUR-ASW** | 3.53% | 3.01 % | 0.95% | 0.56% |
| **CEU-CLM** | 1.40% | 1.16 % | 0.75% | 0.53% |
| **CEU-MXL** | 2.45% | 1.86 % | 1.06% | 0.53% |
| **CEU-CLM-CHB** | 4.77% | 4.60 % | 0.51% | 0.35% |
| **CLM-IBS-ASW** | 4.65% | 4.19% | 0.52% | 0.31% |
| **ACB-CHB-CEU** | 9.00% | 8.87% | 0.36% | 0.34% |

* Leading eigenvalue of the PCA
** Leading eigenvalue of the within-population matrix
*** RMT approximation for the leading eigenvalue of the within-population matrix
**IBS**: Iberian ($n = 147$), **CHB**: Han Chinese in Beijing ($n = 100$), **YRI**: Yoruba ($n = 158$), **CEU**: Utah residents with European ancestry ($n = 104$). **CLM**: Colombians from Medellin Colombia ($n = 102$), **ASW**: Americans of African Ancestry in SW USA ($n = 97$), **PUR**: Puerto Ricans from Puerto Rico ($n = 94$), **MXL**: Individuals of Mexican Ancestry from Los Angeles USA ($n = 100$), **ACB**: African Caribbeans in Barbados ($n = 98$).

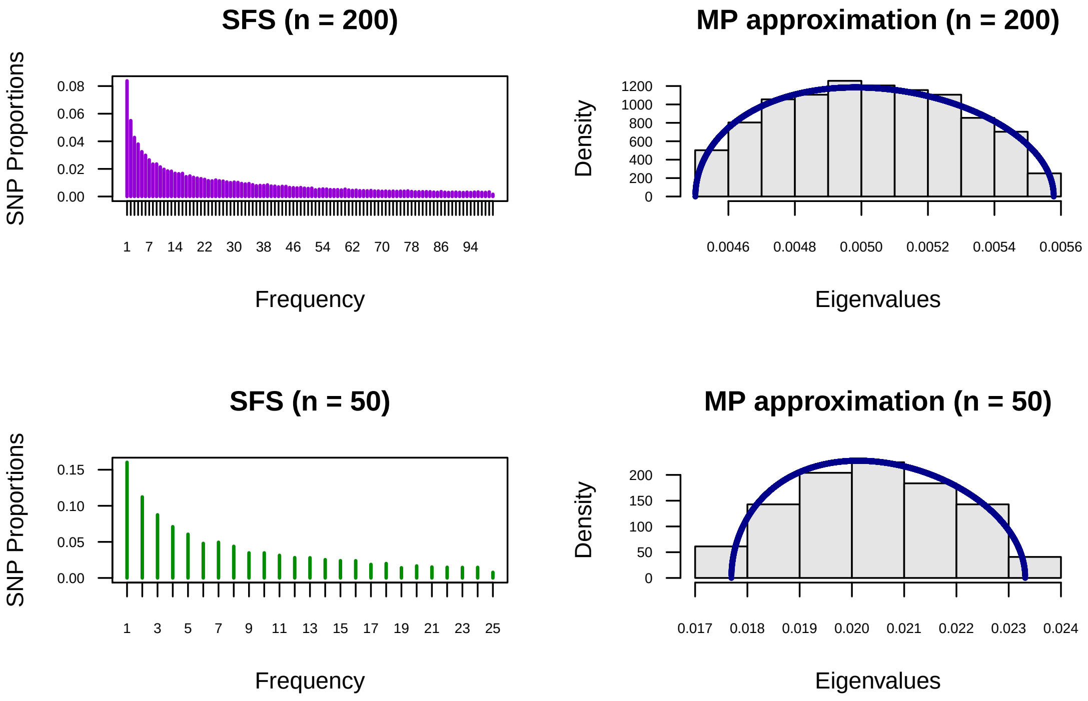

**Figure S1. Random matrix theory approximation of proportions of variance in single population $F$-models.** Two simulations of $F$-models were performed with $p_{\text{anc}}$ drawn from a beta distribution with shape parameters $a = 1$ and $b = 9$, and $F = 15\%$. **Top row**: $n = 200$ individuals and $L = 69,248$ SNPs. **Bottom row**: $n = 50$ individuals and $L = 10,331$ SNPs. **SFS**: Site Frequency Spectrum, **MP approximation**: Marchenko-Pastur approximation of the distribution of scaled PCA eigenvalues (blue curve). Histograms of scaled PCA eigenvalues representing the proportions of variance explained by the $(n - 1)$ principal axes are displayed in grey color.

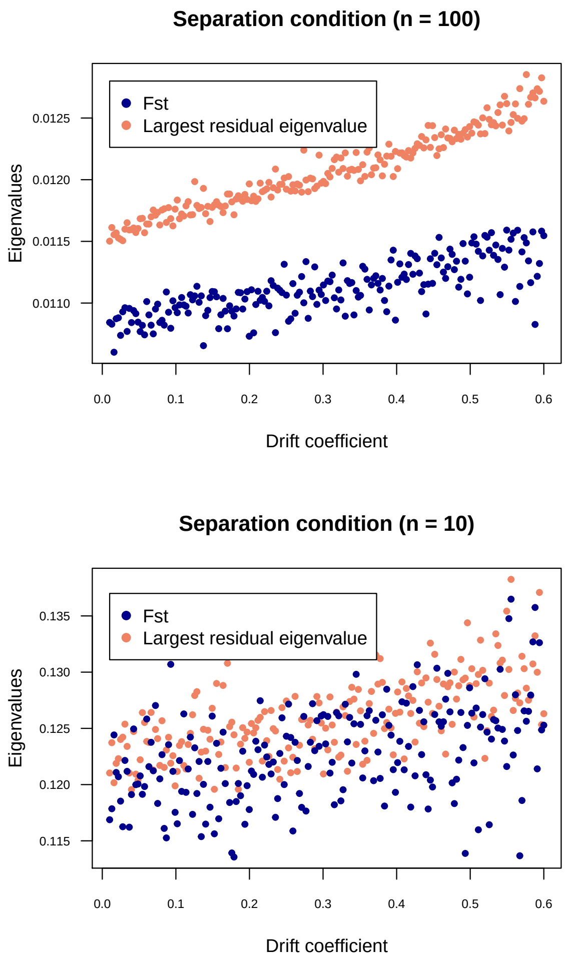

**Figure S2. Separation of variance components in artificial population samples built from single population $F$-models.** For each value of the drift coefficient, each couple of blue and orange dots represent a simulated data set. $F_{\mathrm{ST}}$ (blue dots) corresponds to the non-null eigenvalue of the between-population matrix, $\mathbf{Z}_{\mathrm{ST}}$, and the "residual" value corresponds to the leading eigenvalue of $\mathbf{Z}_{\mathrm{S}}$. Population structure is detected when the blue dot is above the orange dot. **Top row**: 200 simulations with $n = 100$ individuals and $L$ around 10,000 SNPs. **Bottom row**: 200 simulations with $n = 10$ individuals and $L$ around 1,000 SNPs. Simulations were performed with $p_{\mathrm{anc}}$ drawn from a beta distribution with shape parameters $a = 1$ and $b = 9$.

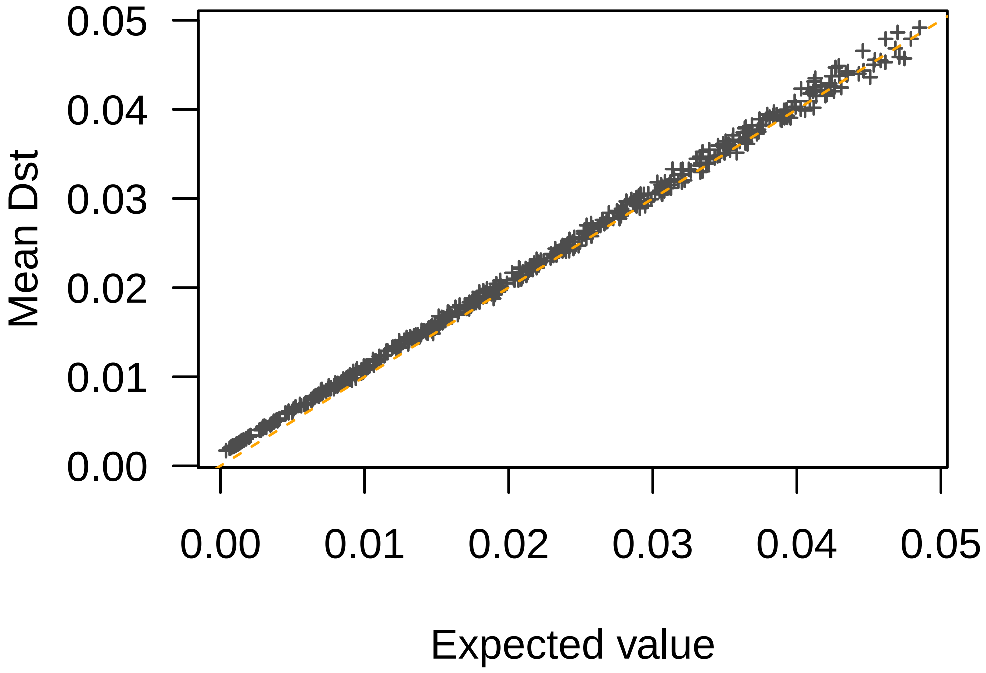

**Figure S3. Accuracy of $D_{\mathrm{ST}}$ estimates with respect to their theoretical values in two-population models.** Comparison of values of $D_{\mathrm{ST}}$ averaged over loci and their theoretical values in $F$-models. Simulations of $F$-models were performed with equal drift coefficients ($F_1 = F_2$) ranging between 1% and 75%, and sample proportions $c_1$ between 10% and 50% ($n = 100$ individuals). The ancestral frequencies, $p_{\mathrm{anc}}$, were drawn from a beta distribution with shape parameters $a = 1$ and $b = 4$.

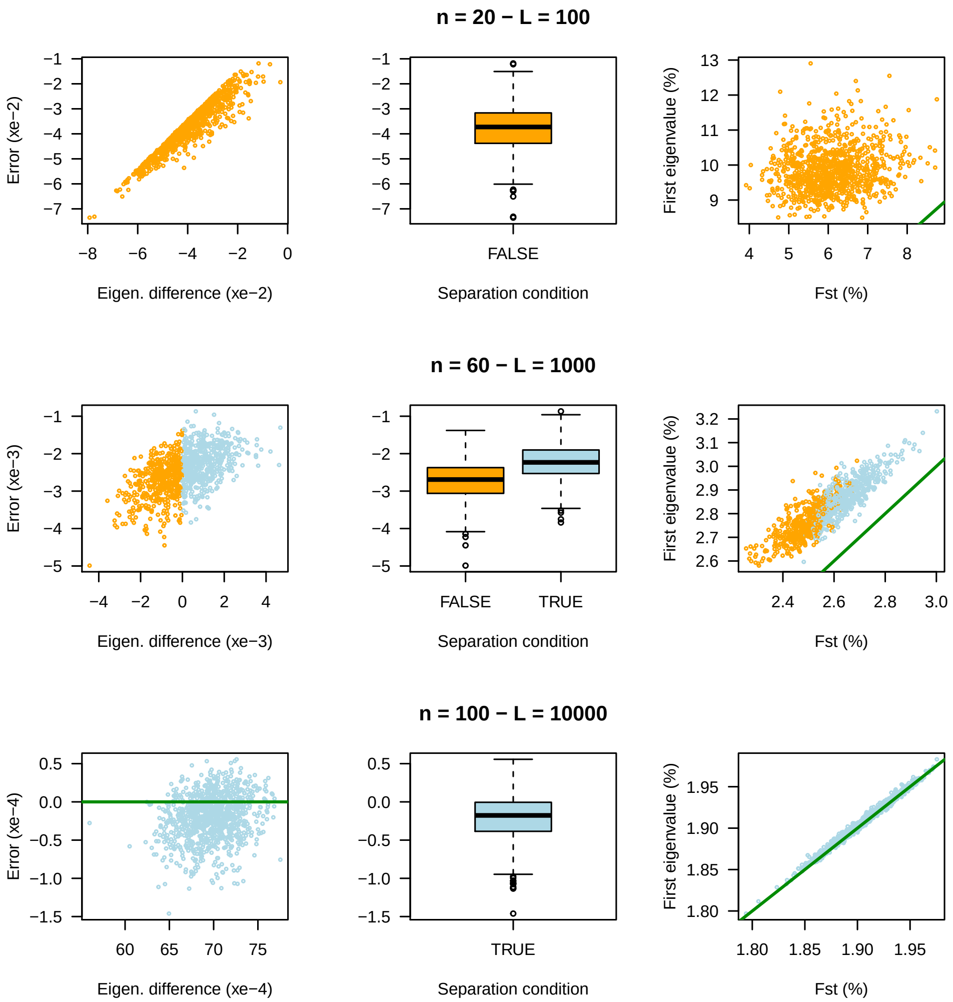

**Figure S4. Accuracy of approximation of $F_{\mathrm{ST}}$ and separation condition in two-population $F$-models. First column**: Approximation error defined as the difference between $F_{\mathrm{ST}}$ and the leading eigenvalue of scaled PCA, $\mathbb{E}[F_{\mathrm{ST}}] - \rho_1^2(\mathbf{Z}^{\mathrm{sc}})/L$, as a function of the difference of eigenvalues, $\rho_1^2(\mathbf{Z}^{\mathrm{sc}}_{\mathrm{ST}})/L - \rho_1^2(\mathbf{Z}^{\mathrm{sc}}_{\mathrm{S}})/L$. **Second column**: Approximation errors according to whether the separation condition is checked or not. **Third column**: Leading eigenvalue of scaled PCA as a function of $\mathbb{E}[F_{\mathrm{ST}}]$. Simulations of $F$-models were performed for $n$ individuals and $L$ loci with equal drift coefficients $F_1 = F_2 = 0.02$. Ancestral frequencies, $p_{\mathrm{anc}}$, were drawn from a beta distribution with shape parameters $a = 1$ and $b = 4$.

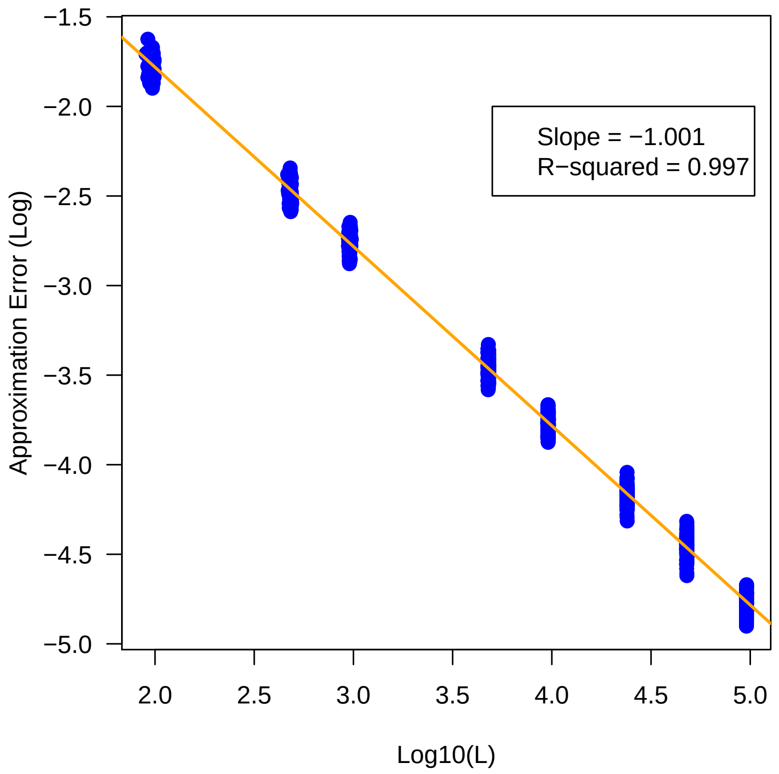

**Figure S5. Approximation errors decrease as $1/L$ in two-population $F$-models.** Approximation error defined as the absolute difference between $F_{\mathrm{ST}}$ and the leading eigenvalue of scaled PCA, $\mathbb{E}[F_{\mathrm{ST}}] - \rho_1^2(\mathbf{Z}^{\mathrm{sc}})/L$ as a function of $1/L$ ($L$ is the number of unlinked loci). The red line corresponds to the linear regression Log(Approx) $= a + b$ Log($L$), and has slope equal to $b = -1.001$ ($R^2 = 0.997$, $P <$ 2e-16). Simulations of $F$-models were performed for $n = 150$ individuals with drift coefficients equal to $F_1 = F_2 = 0.02$. The ancestral frequencies, $p_{\mathrm{anc}}$, were drawn from a beta distribution with shape parameters $a = 1$ and $b = 4$.

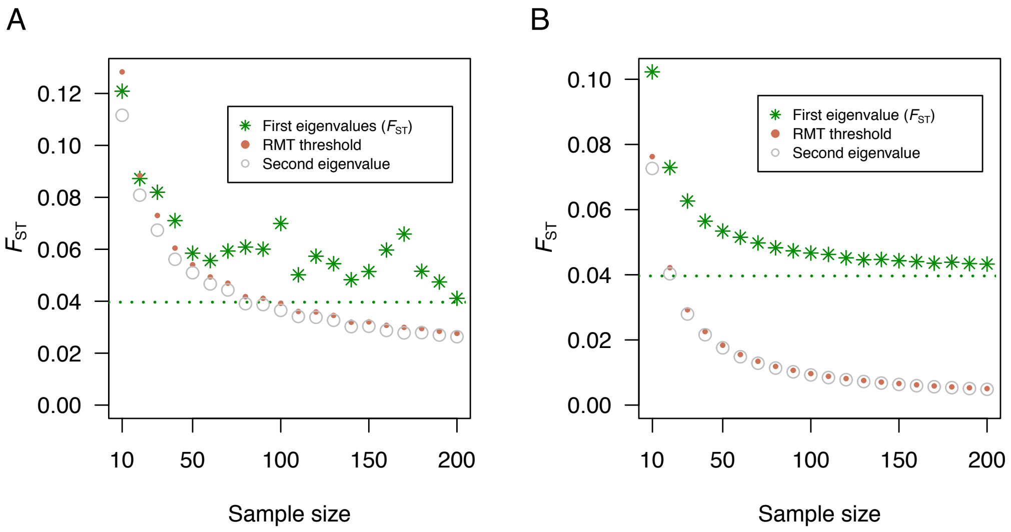

**Figure S6. First eigenvalue ($F_{\mathrm{ST}}$ estimate) as a function of sample size in two-population models.** (**A**) $L = 100$ loci: The separation condition was verified for sample sizes $> 60$. (**B**) $L = 100,000$ loci: The separation condition was verified for all sample sizes. $F_{\mathrm{ST}}$: Leading eigenvalue of the PCA. **RMT threshold**: Approximation of the detection threshold from RMT, equal to $(1/\sqrt{L} + 1/\sqrt{n-1})^2$. Dashed line: Theoretical value for an infinite sample size, $\mathbb{E}[F_{\mathrm{ST}}] = 3.97\%$. Simulations of $F$-models were performed with ancestral frequencies drawn from a beta(1,4) distribution and with $F_1 = F_2 = 10\%$.

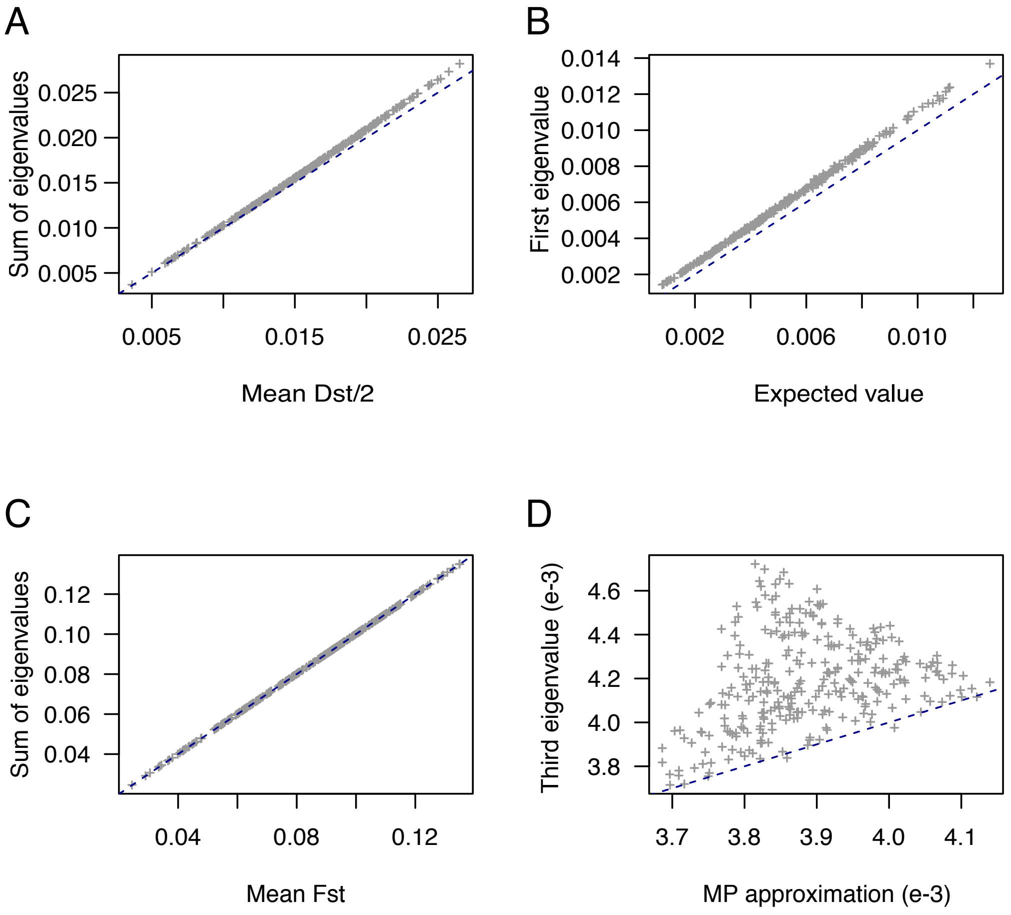

**Figure S7. Leading eigenvalues in three-population $F$-models.** (**A**) Sum of the first two eigenvalues of centered PCA as a function of the mean of $D_{\mathrm{ST}}/2$ across loci. (**B**) First eigenvalue of centered PCA as a function of its expected value $\lambda_1 = (F_1 + F_2 + F_3 + \sqrt{F_1^2 + F_2^2 + F_3^2 - F_1F_2 - F_2F_3 - F_3F_1})/54$. (**C**) Sum of the first two eigenvalues of scaled PCA as a function of the mean of $F_{\mathrm{ST}}$ across loci. (**D**) Third eigenvalue of scaled PCA as a function of its approximation from RMT. **MP approximation**: Marchenko-Pastur approximation of the largest eigenvalue of the residual matrice, $\mathbf{Z}_{\mathrm{S}}/\sqrt{n-3}$, equal to $(1-\rho_1^2-\rho_2^2)\times(1/\sqrt{L}+1/\sqrt{n-3})^2$. The dashed lines correspond to the diagonal $y = x$. Simulations of $F$-models were performed for $n = 100$ individuals with drift coefficients $F_1, F_2, F_3$ between 1% and 25%, equally sampled populations, and ancestral frequencies drawn from the uniform distribution ($L = 20000$ loci).

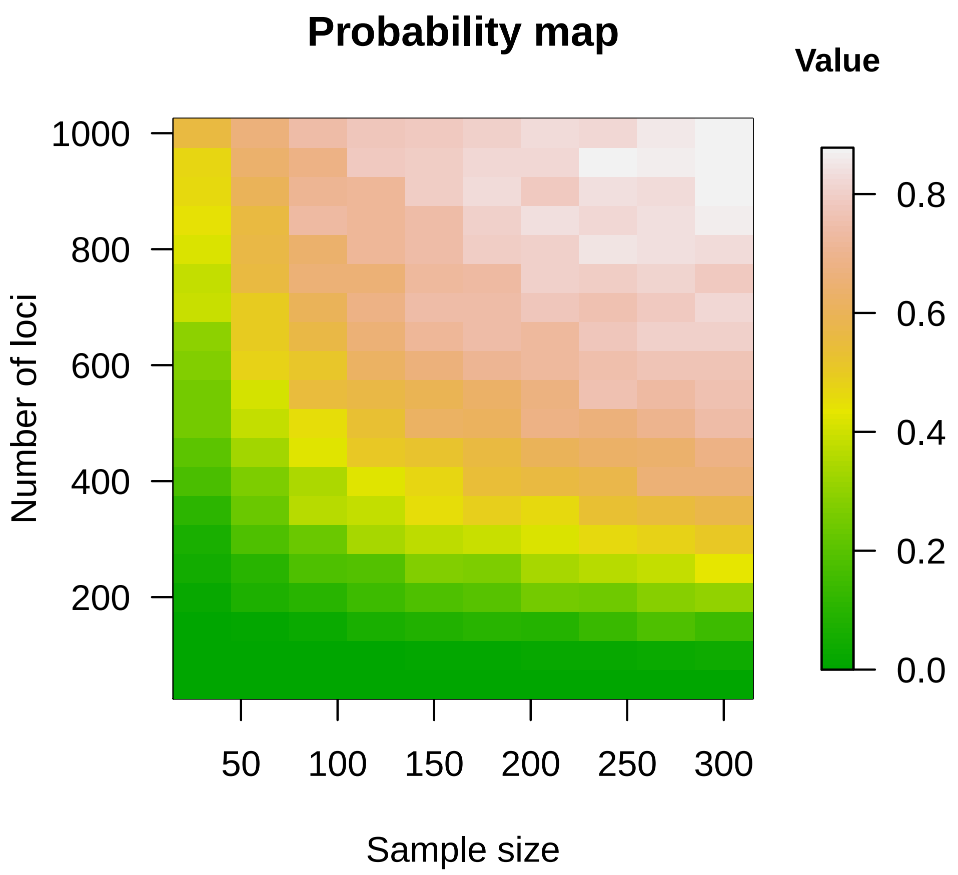

**Figure S8. Separation condition in three-population models.** Probability that the separation condition is verified for sample sizes ranging between $n = 30$ and $n = 300$ individuals, and number of loci ranging between $L = 100$ and $L = 1000$. Simulations of $F$-models were performed with equal sample sizes, random drift coefficients lower than 10%, and ancestral frequencies drawn from the uniform distribution. The number of replicates for each combination of $n$ and $L$ was 500.

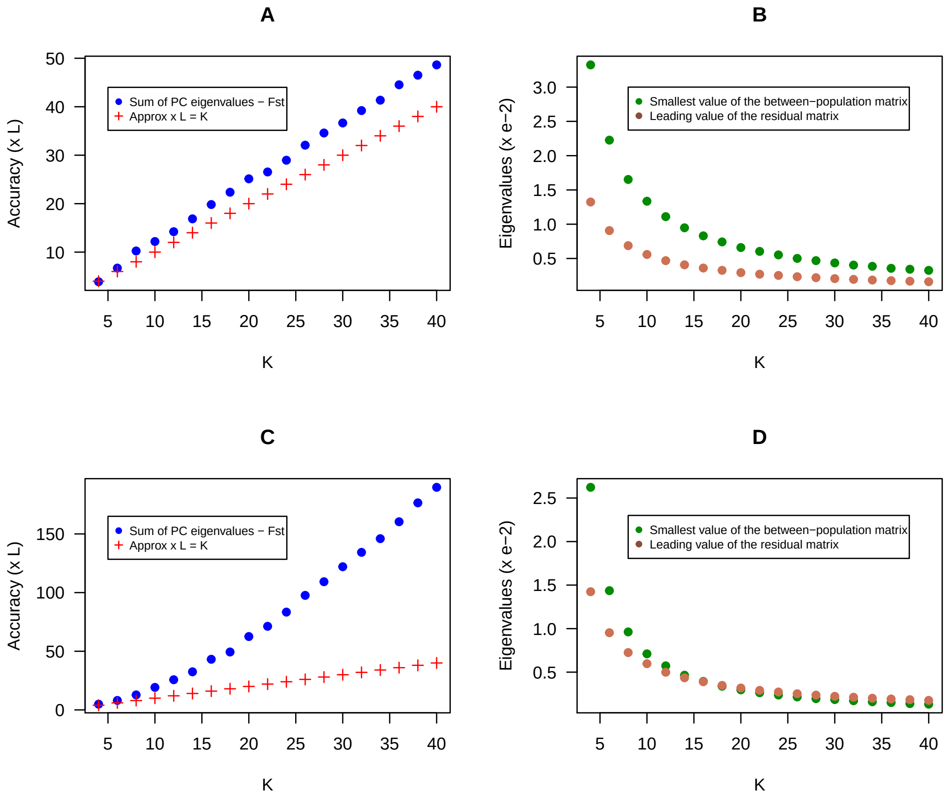

**Figure S9. Accuracy of approximation of $F_{\mathrm{ST}}$ by PCA for $K$-population $F$-models. A-B)** Simulations with equal drift coefficients $F_k = 0.1$. The accuracy of the approximation of $\mathbb{E}[F_{\mathrm{ST}}]$ by the sum of the $K-1$ leading eigenvalues of the PCA is comparable to $K/L$ (A) and the separation of eigenvalues from the residual matrix is verified (B). **C-D)** Simulations with unequal drift coefficients $F_k = 0.2/k$. The accuracy of the approximation of $\mathbb{E}[F_{\mathrm{ST}}]$ diverged from $K/L$ when the separation of eigenvalues did not hold. In all simulations, ancestral frequencies were drawn from a beta distribution with shape parameters $a = 1$ and $b = 4$. Sub-population samples had size equal to $n_k = 20$, the total population size was $n = 20 \times K$), and the number of SNP loci was $L \approx 19850$.

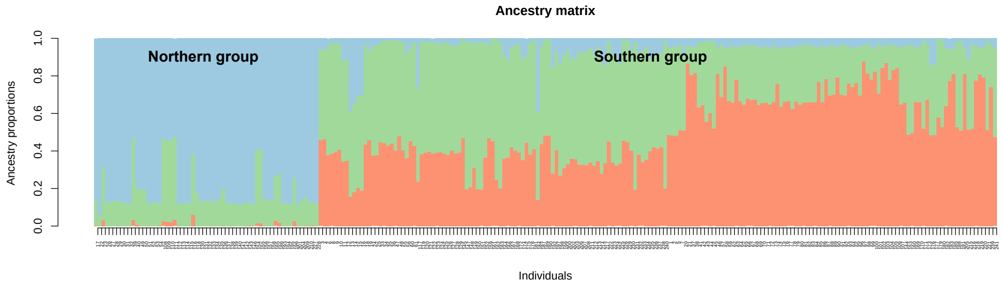

**Figure S10. Estimates of ancestry coefficients for 241 Swedish accessions of *A. thaliana*.** Ancestry coefficients obtained from the spatially explicit ancestry estimation program `tess3r` with $K = 3$ populations. The southern group exhibits substantial levels of mixed ancestry.
